# HSC70 prevents TDP-43 nuclear puncta formation and toxicity in a novel nuclear puncta cell model for ALS

**DOI:** 10.64898/2026.07.21.739729

**Authors:** Jeng Jung Wu, Shin-Yu Fan, Tien-Hsien Chang, Yun-Ru Chen

## Abstract

Amyotrophic lateral sclerosis (ALS) is categorized by TDP-43 proteinopathy, however, the nuclear pathological events remain poorly defined. While cytoplasmic TDP-43 inclusions dominate the late disease stages, accumulating evidence indicates that nuclear TDP-43 assemblies arise earlier and impair RNA splicing. Here, we characterized a single RRM-proximal TDP-43 variant, G148V, designed to disrupt nucleic-acid engagement without altering canonical RNA-binding residues. Structural and biophysical analyses revealed conformational changes and loss of DNA/RNA binding. In mammalian cells, TDP-43 G148V robustly formed nuclear puncta with high penetrance, exhibiting solid-like properties, pathological phosphorylation, splicing dysfunction, and toxicity. Furthermore, we identified molecular chaperone HSC70 as an important regulator of the nuclear puncta assembly. HSC70 redistributed into G148V nuclear puncta to modulate their material state, whereas HSC70 depletion significantly promoted puncta solidification, increased insoluble TDP-43 accumulation, and enhanced cytotoxicity. Disease-associated K181E and K263E mutants also formed nuclear puncta and induced HSC70 nuclear redistribution. These findings establish G148V as a model of early nuclear TDP-43 pathology and highlight HSC70-mediated regulation as a key factor of TDP-43 nuclear assembly.

**Highlights:**

- A single TDP-43 mutation, G148V, in RRM1 domain robustly induces nuclear puncta without exogenous stress.
- G148V disrupts nucleic-acid binding, driving solid-like nuclear assemblies with hyperphosphorylation.
- Nuclear G148V puncta impair splicing regulation and reduce cell viability, recapitulating early ALS pathology.
- The molecular chaperone HSC70 modulates puncta material states and mitigates G148V-associated cytotoxicity.

**Graphical abstrac:** 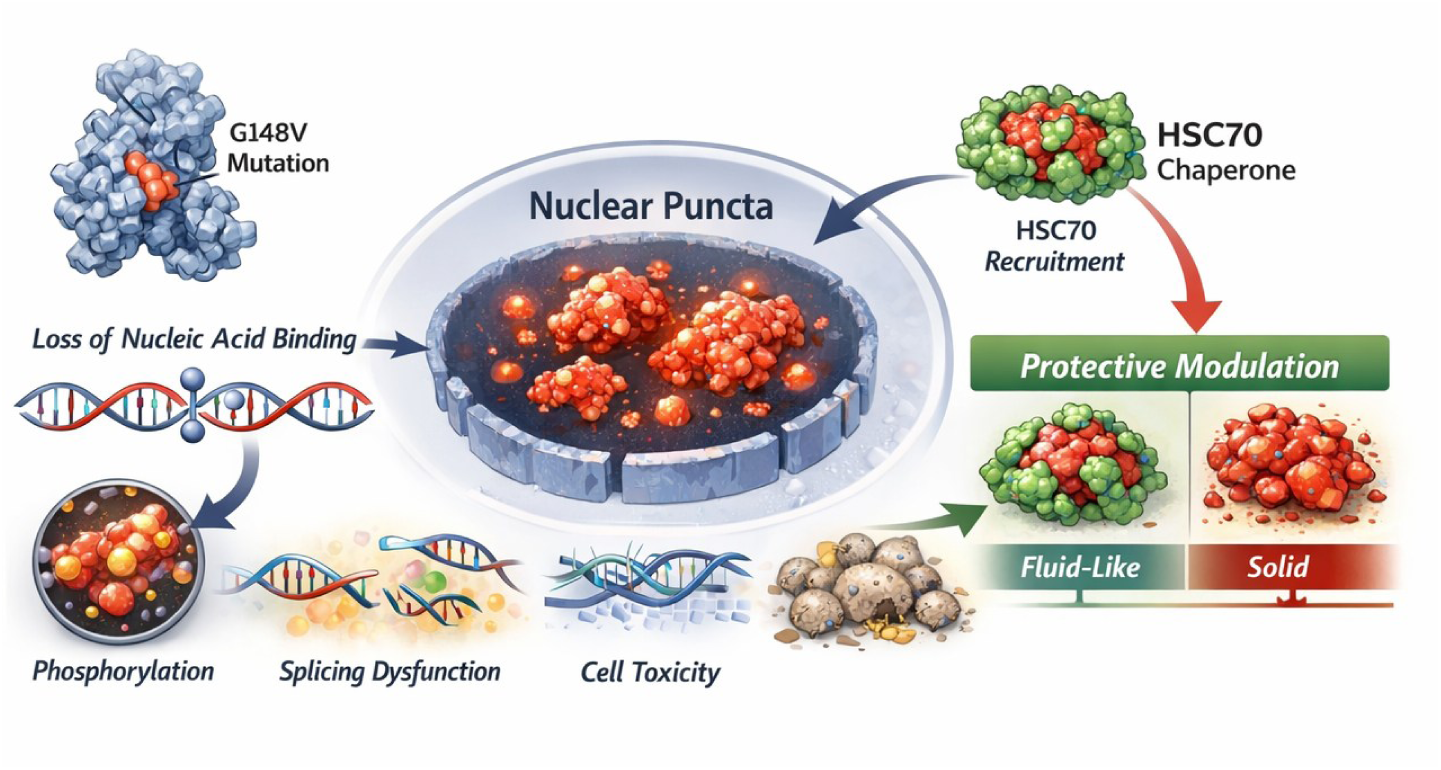

*eTOC blurb:* A structure-guided TDP-43 G148V mutation reveals how loss of nucleic-acid engagement promotes early nuclear condensation, splicing dysfunction, and toxicity, while uncovering a protective role for HSC70 in regulating condensate properties during ALS pathogenesis.

## Introduction

Amyotrophic lateral sclerosis (ALS) is a relentless progressive neurodegenerative disorder marked by the selective loss of motor neurons in the cortex, brainstem, and spinal cord, leading to muscle weakness, spasticity, and ultimately respiratory failure (Brown & Al-Chalabi, 2017, Hardiman, Al-Chalabi et al., 2017, Kiernan, Vucic et al., 2011, Nijs & Van Damme, 2024, Yuan, Yang et al., 2025). Approximately 10% of ALS cases are familial, while the majority are sporadic, yet both converge on overlapping molecular pathways (Mengistu et al., 2025, Nijs & Van Damme, 2024, Taylor, Brown et al., 2016, Yuan et al., 2025). Despite decades of research, ALS therapies remain limited, with Riluzole and Edaravone providing only modest benefit and newer approaches such as antisense oligonucleotides showing incremental but still insufficient progress (Boros, Schoch et al., 2022, Hayes & Kalab, 2022, Jaiswal, 2019).

A central molecular hallmark of ALS is dysfunction of TAR DNA-binding protein 43 (TDP-43), encoded by the TARDBP gene (Feneberg, Thompson et al., 2025, Neumann, Sampathu et al., 2006, Prasad, Bharathi et al., 2019). TDP-43 is a 414-amino-acid protein composed of an N-terminal domain, a nuclear localization signal, two RNA recognition motifs (RRM1 and RRM2), a nuclear export signal, and a disordered C-terminal domain (Figure 1A). Under normal conditions, TDP-43 resides in the nucleus and regulates transcription, splicing, RNA transport, and stability (Carmen-Orozco, Tsao et al., 2024, Decker, Menge et al., 2025, Prasad et al., 2019). In nearly all sporadic ALS cases and many familial forms, TDP-43 is depleted from the nucleus and accumulates in the cytoplasm, where it becomes ubiquitinated and hyperphosphorylated (Neumann et al., 2006). These inclusions contribute to neuronal dysfunction through both loss-of-nuclear function and toxic gain-of-function in the cytoplasm (2025, Taylor et al., 2016). More recently, nuclear TDP-43 pathology has gained increasing attention besides cytoplasmic inclusions (Carmen-Orozco et al., 2024, Decker et al., 2025, Huang, Ellis et al., 2024, Spence, Waldron et al., 2024). Normally, TDP-43 forms dynamic nuclear condensates via liquid-liquid phase separation to compartmentalize RNA processing (Carmen-Orozco et al., 2024). Perturbations such as mutations, post-translational modifications, or chronic stress can destabilize this equilibrium, causing nuclear puncta to harden into insoluble inclusions (Feneberg et al., 2025, Keating, Bademosi et al., 2023, Yabata, Riku et al., 2023). These aggregates sequester functional TDP-43 and other RNA-binding proteins, leading to cryptic exon inclusion, splicing errors, and transcriptome disruption (Decker et al., 2025, Huang et al., 2024). Importantly, recent work using RNA aptamers has revealed that nuclear TDP-43 aggregation represents an early pathological event coinciding with STMN2 cryptic splicing defects (Spence et al., 2024). Evidence from cellular and animal models further supports that nuclear puncta may represent an early stage in the pathological cascade, preceding cytoplasmic inclusions observed in patient tissue (Feneberg et al., 2025, Yabata et al., 2023). However, current cellular systems have struggled to reliably reproduce nuclear TDP-43 puncta under physiological conditions. Most models require chemical stressors or multi-site TDP-43 mutants, while single-point variants typically induce puncta at low efficiency (Huang et al., 2024, Keating et al., 2023, Prasad et al., 2019).

**Figure 1.**
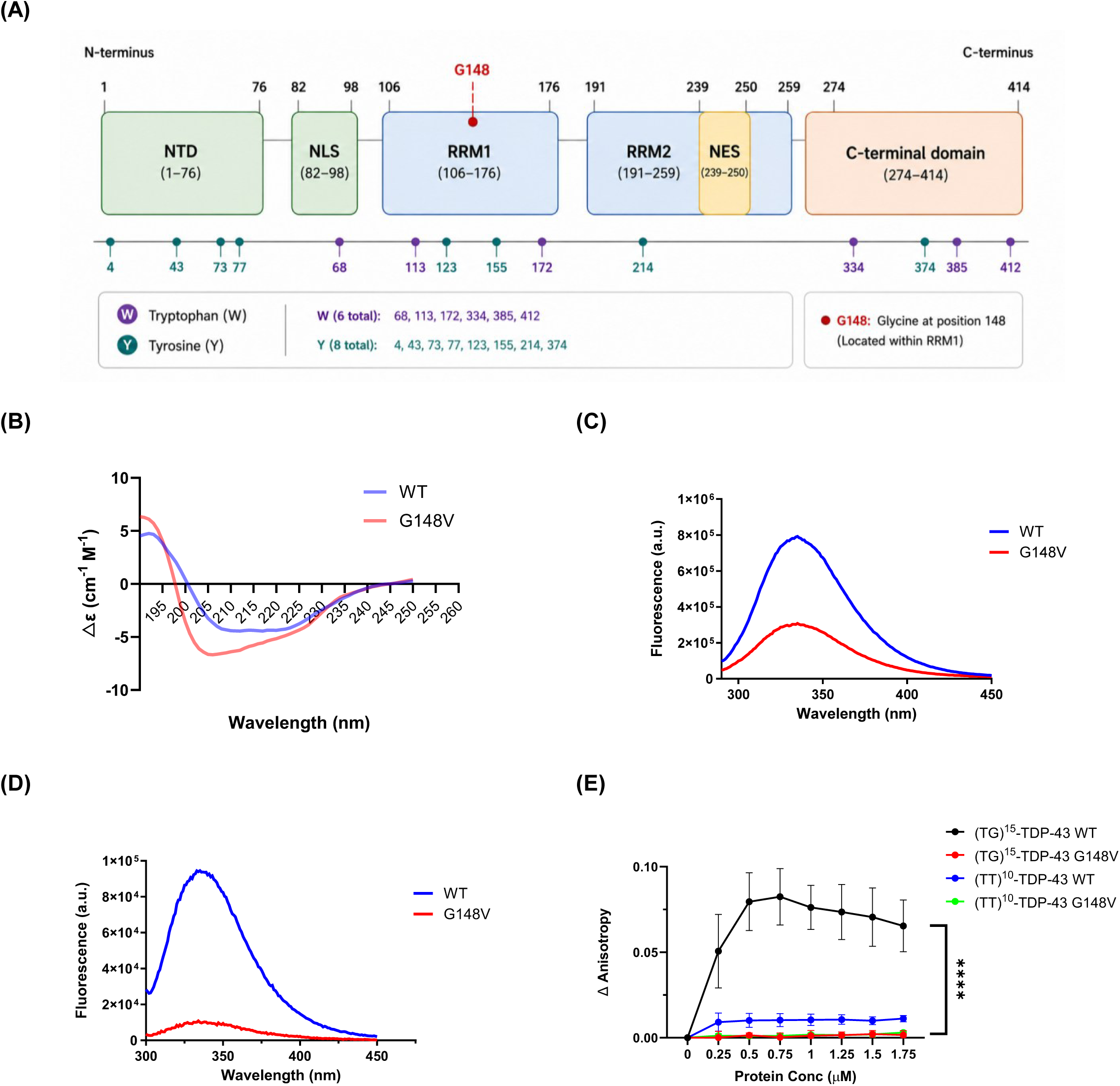
The conformation and DNA binding of recombinant TDP-43 G148V were different from WT. (A) Schematic representation of TDP-43. NTD, N-terminal domain; NLS, nuclear localization signal; RRM, RNA recognition motif; NES, nuclear export signal. (B) Far-UV circular dichroism demonstrated that WT and G148V TDP-43 both showed α-helical and β-strand structures, but G148V exhibited altered ellipticity. (C) Intrinsic fluorescent spectra of 280 nm indicated that G148V exhibited reduced fluorescence, indicating lower solvent exposure than WT. (D) Intrinsic fluorescent spectra of 295 nm indicated that tryptophan of G148V exhibited reduced fluorescence. (E) The result of anisotropy assay. G148V showed no binding to either probe, indicating impaired nucleotide binding. (n = 3 biological replicates) Data information: All data are presented as mean ± SD. In (F), TT, a T-repeated oligomer probe as the negative control. TG, a TG repeated oligomer probe, was used as the positive control. ****P < 0.0001 (Two-way Anova).

In this context, molecular chaperones such as heat shock cognate protein 70 (HSC70, HSPA8) have emerged as important modulators of ALS pathology (Arosio, Cristofani et al., 2020, Coyne, Lorenzini et al., 2017, Huang, Bose et al., 2014). HSC70 is a constitutively expressed member of the HSP70 family that maintains proteostasis by assisting in protein folding, refolding denatured proteins, and targeting misfolded proteins for degradation (Kalmar, Lu et al., 2014, Pustovaya, Venediktov et al., 2026, Stricher, Macri et al., 2013). In neurons, HSC70 is essential for sustaining protein quality control under high metabolic demand (Coyne et al., 2017). In ALS, HSC70 directly interacts with TDP-43 and. (Coyne et al., 2017, Huang, Bose et al., 2014). Experimental models have shown that mutant TDP-43 can reduce synaptic HSC70 levels, impairing synaptic vesicle cycling and exacerbating neurodegeneration Coyne et al., 2017). Conversely, HSC70 overexpression partially rescues TDP-43-mediated toxicity and restores synaptic function in experimental ALS models, highlighting its neuroprotective potential (Coyne et al., 2017).

In this paper, we report a unique TDP-43 mutation G148V, which alters the nucleic acid binding motif, consistently generates nuclear puncta in cultured cells. These puncta exhibit hallmark features of TDP-43 proteinopathy, including solid-like aggregation, insoluble products, hyperphosphorylation, cytotoxicity, and splicing abnormalities. Interestingly, the G148V mutant uncovered a functional link between TDP-43 and HSC70. During puncta formation, HSC70 undergoes a marked redistribution from the cytoplasm into the nucleus, where it colocalizes with nuclear TDP-43 assemblies. Knockdown experiments revealed that HSC70 modulates both the fluidity and phosphorylation state of these puncta, indicating a regulatory role in their biophysical and biochemical properties. Comparable HSC70 mislocalization was also observed in disease-associated TDP-43 mutants. However, the G148V variant displayed greater stability and reproducibility in puncta formation, making it a valuable tool for mechanistic studies. Collectively, our findings identified HSC70 as a critical factor of nuclear TDP-43 puncta dynamics and established the G148V mutant as a promising cell-based model for early ALS pathogenesis. This system provides a reproducible platform for dissecting the molecular mechanisms underlying nuclear TDP-43 pathology and offers new opportunities for therapeutic exploration.

## Result

### The G148V mutation alters the structure and characteristics of TDP-43

Previous crystallographic studies have identified residues F147 and F149 within RRM1 of TDP-43 are the critical residues for nucleic acid binding (Kuo, Chiang et al., 2014). G148 is positioned between F147 and F149 and is a glycine residue that contributes to peptide flexibility. Hence, we hypothesized that altering this residue could modulate nucleic acid-binding properties without directly disrupting the canonical binding residues. To understand the possibly interruption of nucleic acid binding in TDP-43, we generated a mutation at residue 148 from glycine to valine (G148V) to increase the hydrophobicity and enlarge the side chain. Full-length TDP-43 recombinant protein with or without G148V mutation were expressed and purified from E. coli. The purified recombinant proteins were subjected to structural analyses to evaluate the conformational changes of the single-residue substitution. The TDP-43 wild-type (WT) and G148V proteins were subjected to far-ultraviolet circular dichroism (far-UV CD) spectroscopy for detecting the secondary structures and intrinsic fluorescence spectroscopy for monitoring the tertiary structural changes. The far-UV CD spectra showed that both WT and G148V TDP-43 exhibited some degree of α-helical and β-strand structures, the ellipticity of the G148V mutation was different from that of WT TDP-43 (Fig. 1B). The structure prediction using software K2D3 revealed that the WT and G148V contained α-helical content of 33.01% and 37.24%, and β-strand content of 17.82% and 12.63%, respectively, suggesting that the G148V mutation increase the α-helical content while decreasing the β-strand content of TDP-43. The intrinsic fluorescence spectra excited at 280 and 295 nm showed similar emission spectra at ∼335 nm for both proteins, however, the fluorescence intensity was lower in G148V indicating the potential quenching of tyrosine and tryptophan residues in the G148V mutant compared to WT, indicating that the G148V mutation is less solvent-exposed (Fig. 1C,D). These suggested that one single mutant glycine to valine might change the conformation of TDP-43.

Given the dysfunction of the associated structural changes, we investigated whether this mutation affects the DNA- and RNA-binding capacity of TDP-43. For this purpose, we employed the anisotropy assay by titrating TDP-43 to the Alexa488 fluorescently labeled single-strand DNA. The binding of the fluorescently labeled nucleotides to TDP-43 will result in an increase of the anisotropy signal. The DNA with TG repeats that previously known to bind TDP-43 served as a positive control, whereas the single-strand DNA TT repeat that did not bind TDP-43 served as a negative control (Chang, Wu et al., 2012, Kuo et al., 2014). The result showed that the anisotropy signal for WT titrating to the TG repeat gradually increased upon rising TDP-43 concentration, whereas it exhibited a low and much weaker binding ability to the TT probe. In contrast, the signal of the G148V mutant didn’t show binding to both probes demonstrating the loss of its nucleotide binding ability (Fig. 1D). The structural alterations and the loss of nucleic acid-binding capacity observed in the G148V mutant indicate that this single-residue substitution is sufficient to disrupt an essential functional property of TDP-43 while leaving the known binding residues F147 and F149 intact. These findings suggest that the G148V variant could be used as a model for functional impairment consequences relevant to TDP-43 pathology.

### TDP-43 G148V formed hyperphosphorylated and solid-like puncta in the nucleus in cell

Next, to understand the G148V mutation effect in cell, we expressed WT TDP-43 and G148V variant to human embryonic kidney cell HEK293T (Fig. 2A) and mouse neuroblastoma N2a cell lines (Appendix Fig. S1A). Interestingly, while both WT and G148 TDP-43 expressed in the nucleus, the G148V mutation accumulated into numerous small puncta while WT diffused throughout the nucleus (Fig. 2A; Appendix Fig. S1A). Quantifying the cells containing puncta demonstrated that the G148V mutation produced significantly smaller puncta in the nucleus than WT (Fig. 2B; Appendix Fig. S1B). Also, the cross-section analysis of the fluorescence profile of the G148V mutation was more heterogenicity compared to WT, demonstrating G148V mutation accumulated into small puncta in the nucleus (Fig. 2C; Appendix Fig. S1C). In addition, to exclude the potential influence of the fusion protein, TDP-43 G148V carrying only a FLAG tag was also included in the analysis, and similar results were observed (Appendix Fig. S2A,B).

**Figure 2.**
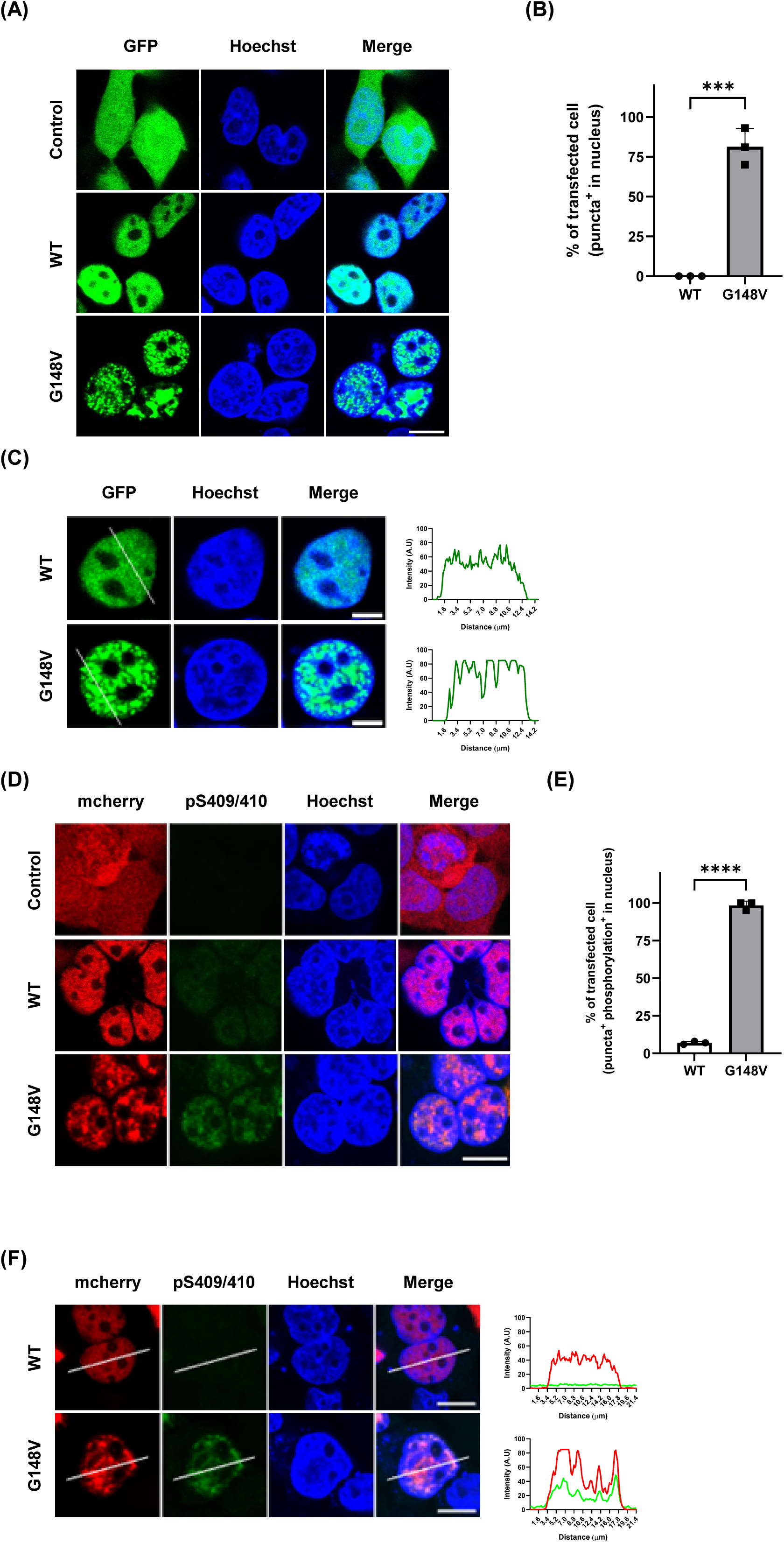
TDP-43 G148V accumulates as puncta in the nucleus in HEK293T cell. (A) TDP-43 G148V formed nuclear puncta while WT TDP-43 showed a diffused expression in HEK293T. Scale bar: 20 μm. (B) Quantitative result of the transfected cells with the nuclear puncta (n = 3 biological replicates). (C) Intensity analysis of the nuclear TDP-43 expression. The intensity peaks of G148V are more heterogenicity than WT in the nucleus. Scale bar: 5 μm. (D) Phosphorylated TDP-43 increased in the TDP-43 G148V puncta of HEK293T nucleus. Scale bar: 5 μm. (E) Quantitative result of the phosphorylated TDP-43 level (n = 3 biological replicates). (F) Intensity analysis of the nuclear TDP-43 expression. The intensity peaks of G148V and pS409/S410 overlapped in the nucleus. Scale bar: 10 μm. Data information: All data are presented as mean ± SD. In (B)(E), n = ∼80 cells, ***P = 0.0003, ****P<0.0001 (Student’s t-test).

Aberrant hyperphosphorylation of TDP-43, particularly at residues S409/410, represents a hallmark of ALS pathology (de Boer, Orie et al., 2020, Kellett, Bademosi et al., 2025). Here, we investigate whether the nuclear puncta formed by the G148V mutation exhibits such characteristics by immunofluorescence probed by the hyper-phosphorylation sites S409 and S410-specific antibody. The result showed that the phosphor-TDP-43 signal almost completely co-localized with the puncta signal in G148V in both HEK293 (Fig. 2D) and N2a cells (Appendix Fig. S1D), whereas WT did not show hyperphosphorylated TDP-43 signal. Quantitative analysis revealed that cells expressing the G148V mutation were more prone to forming highly phosphorylated puncta (Fig. 2E; Appendix Fig. S1E). Furthermore, the cross-section analysis of fluorescence intensity profiling demonstrated that the profile of the G148V mutation closely corresponded with the phosphorylation profile in both cells (Fig. 2F; Appendix Fig. S1F). TDP-43 G148V carrying only a FLAG tag was also observed similar results (Appendix Fig. S2C). According to the literature, since many known kinases of TDP-43 are active in both the nucleus and cytoplasm and have been observed to be upregulated in the patients, it is inferred that TDP-43 may undergo hyperphosphorylation in the nucleus at an early stage (Kellett et al., 2025, Nilaver & Urbanski, 2023). This indicates that when TDP-43 forms abnormal puncta within the cell nucleus, it exhibits a highly phosphorylated pathological characteristic.

Previous reports have showed that TDP-43 is capable of undergoing liquid-liquid phase separation (LLPS), giving rise to dynamic condensates in cell (Liu, Xiang et al., 2025, Song, 2024).Therefore, we investigated whether the small puncta formed by TDP-43 G148V mutation exhibit LLPS properties by the fluorescence recovery after photobleaching (FRAP) assay (Fig. 3A). The mCherry-fused TDP-43 WT or G148V was transiently expressed in HEK293T cells and 20 puncta were selected among 20 cells for FRAP. The quantitative result is shown in Fig 3B and C. The results showed that WT TDP-43 recovered rapidly during post-bleaching to reach 60% of the fluorescence intensity, whereas the small puncta formed by the G148V mutant showed two types of FRAP property. The major group representing ∼80% of the puncta exhibited a recovery efficacy comparable to WT, while the minor group representing ∼20% of the puncta demonstrated a poor recovery indicating a solid-like property. Together, the results showed that G148V mutant forms hyperphosphorylated and solid-like puncta more readily than WT in the nucleus (Fig. 3C), recapitulating disease phenotype with the nuclear TDP-43 accumulation and hyperphosphorylation.

**Figure 3.**
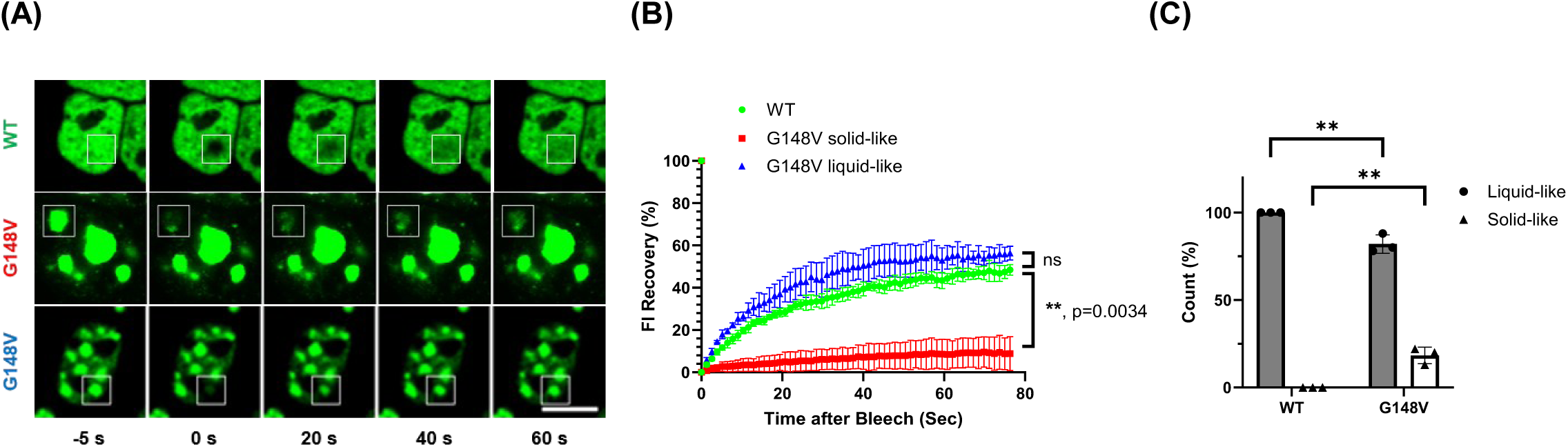
TDP-43 G148V formed solid-like puncta in the nucleus of HEK293T cell. (A) FRAP assay for nuclear puncta. The fluorescence of G148V could be separated into two groups, liquid-like and solid-like. Scale bar: 10 μm. (B) FRAP analysis of the nuclear TDP-43. The trend chart demonstrated a significant difference between WT and solid-like G148V puncta. (C) Quantification of liquid- and solid-like puncta (n = 3 biological replicates). Data information: All data are presented as mean ± SD. In (H), n = ∼20 cells, **P = 0.0034 (Two-way Anova). In (I), n = ∼20 cells, **P = 0.0041 (liquid-like) and 0.0026 (solid-like) (Student’s t-test).

### TDP-43 G148V formed the insoluble fraction that affects cell viability and the splicing process

Previous biochemical fractionation studies from the postmortem tissues or cell line models from ALS/FTD patients demonstrate a shift from soluble TDP-43 to detergent-insoluble TDP-43 aggregates (Kellett et al., 2025, Tsekrekou, Giannakou et al., 2024). To examine whether the G148V mutation shows a higher propensity to form insoluble aggregates, we subjected the cellular fractions expressing mCherry-fused TDP-43 to Western blot. The results showed that the expression level of monomer in the soluble fraction decreased in the G148V variant, whereas the expression level of the insoluble fraction increased (lower red arrow) compared to WT (Fig 4A,B). Notably, the G148V variant displayed enhanced accumulation of detergent-resistant high-molecular-weight (HMW) species (upper red arrow), compared to WT. Similar results were observed in mouse neuroblastoma N2a cells (Appendix Fig. S3A,B).

**Figure 4.**
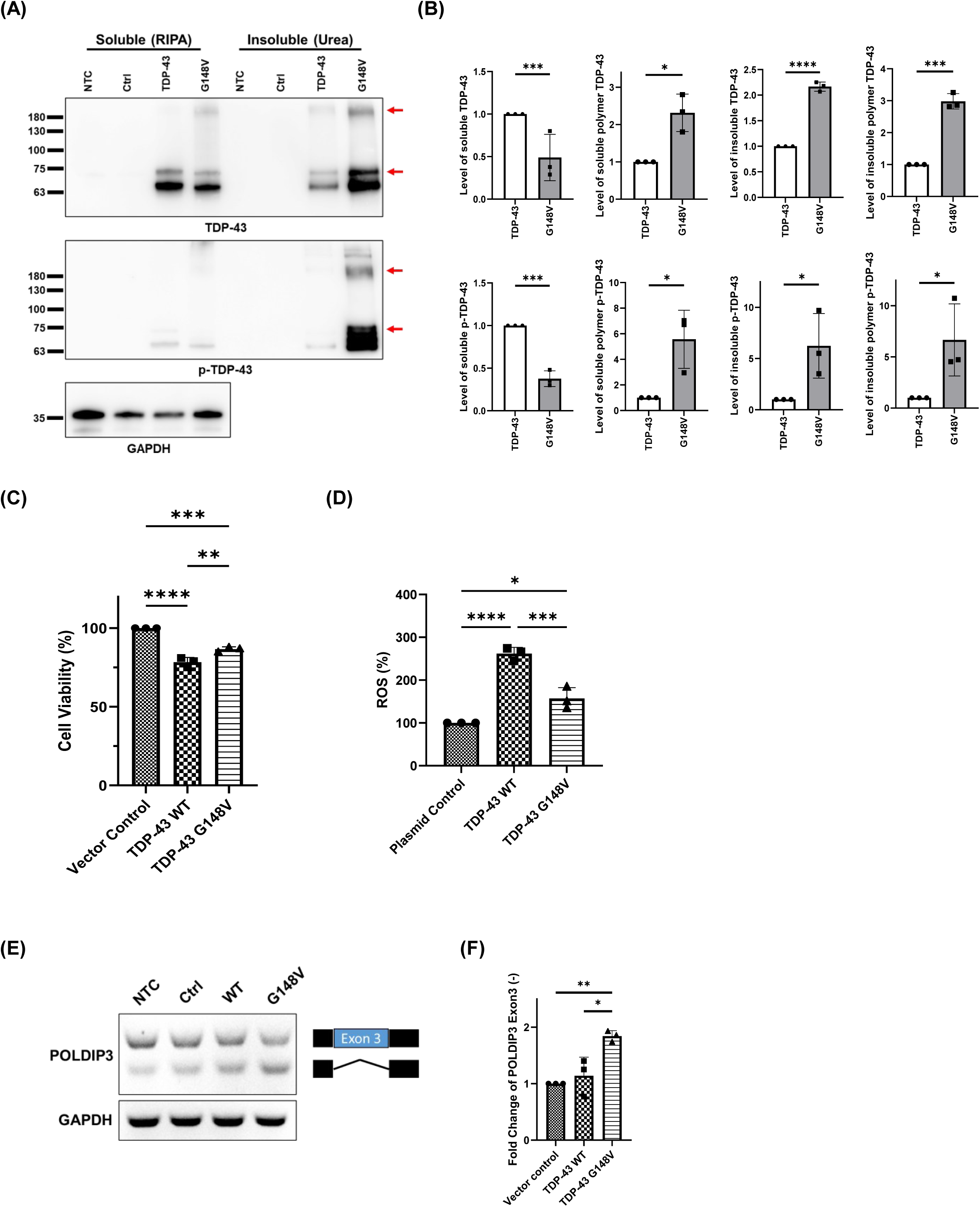
Insoluble fraction formed by G148V mutation, affecting the cell viability and splicing. (A) Expression levels of TDP-43 G148V in HEK293T cells. Soluble protein was extracted with RIPA buffer, and insoluble protein with Urea buffer. (B) Quantitative analysis of the western blot. G148V showed increased insoluble, hyperphosphorylated monomer and SDS-resistant HMW species compared with WT. (n = 3 biological replicates) (C) CCK-8 assays showed reduced viability in HEK293T cells expressing WT or G148V, with G148V showing a nonsignificant trend toward higher viability than WT. (n = 3 biological replicates) (D) The cell viability examined by the ROS assay. WT and G148V TDP-43 both increased ROS relative to the control. The ROS of G148V was lower than WT TDP-43. (n = 3 biological replicates). (E) Representative RT-PCR of POLDOP3 exon 3 alternative splicing. (F) Quantification of POLDIP3 exon 3(−). G148V, but not WT, promoted POLDIP3 exon 3 skipping, indicating disrupted splicing activity. (n = 3 biological replicates) Data information: All data are presented as mean ± SD. In (A), the upper and lower red arrows indicate polymer and monomer, respectively. In (B), *P < 0.05, ***P < 0.001, ****P < 0.0001 (Student’s t-test). In (C), *P = 0.0305, **P = 0.0062 (One-way Anova with Tukey’s multiple comparisons test). In (D), *P = 0.0152, ***P = 0.0007, ****P < 0.0001 (One-way Anova Anova with Tukey’s multiple comparisons test). In (F), *P = 0.0114, **P = 0.0048 (One-way Anova with Tukey’s multiple comparisons test).

Meanwhile, we examined the hyperphosphorylated TDP-43 and found that the insoluble G148V monomer was significantly increased (Fig. 4A,B), while the soluble phospho-TDP-43 was reduced. Meanwhile, the insoluble hyperphosphorylated HMW species also increased in G148V compared to WT. Therefore, G148V mutant demonstrated a higher propensity to form insoluble and hyperphosphorylated monomer as well as SDS-resistant HMW species compared to WT. These findings indicate that the TDP-43 G148V mutant tends to accumulate into SDS-resistant and insoluble aggregates with hyperphosphorylation. The result is consistent with fluorescence imaging and FRAP assay.

Next, we asked whether the cell survival was affected by the property of G148V by cell survival and toxicity assays. To avoid potential confounding effects of the fluorescent protein on TDP-43, cell viability assays were performed using a FLAG-tagged construct. Cell counting kit-8 (CCK-8) and reactive oxygen species (ROS) assay were utilized to measure dehydrogenase activity and ROS in cells to assess their viability. CCK-8 assays showed that cell viability was significantly reduced in the HEK293T cells expressing either WT or G148V compared with the vector control (Fig. 4C). However, the viability was slightly higher in G148V-expressing cells than in those expressing WT. These results indicate that the G148V mutant retains cytotoxicity. In addition, ROS measurements showed that, relative to the control, cells expressing wild-type TDP-43 exhibited a higher degree of ROS; cells expressing the G148V mutant displayed higher ROS levels compared to the vector control, but lower than the wild-type (Fig. 4D). These findings are largely consistent with the CCK-8 results, given that ROS generation is an early event preceding mitochondrial impairment. Although cells expressing the G148V mutant exhibited lower ROS levels than those expressing wild-type TDP-43, the increased ROS production still exerted adverse effects on cellular viability. These findings suggest that nuclear puncta formed by G148V are prone to generate insoluble species that resist cellular degradation, ultimately leading to decreased cell viability. However, the nuclear puncta formation by G148V has less toxicity than WT TDP-43 indicating the solid puncta may ameliorate the toxicity in comparison to the diffused TDP-43 species.

The dysregulation of TDP-43 has been shown to lead to abnormal splicing events known as “cryptic exons.” To determine whether the G148V mutation affects splicing function, the alternative splicing of POLDIP3, an endogenous gene whose exon 3 inclusion is performed by functional TDP-43, was performed in the HEK293T cell line (Chen, Topp et al., 2019, Huang et al., 2024, Shiga, Ishihara et al., 2012). Under normal conditions, exon 3 of POLDIP3 is retained; however, pathological settings have been associated with abnormal skipping of exon 3. Expression of the WT TDP-43 showed minimal difference compared to the control group. However, expression of the TDP-43 G148V mutant resulted in exclusion of exon 3 of POLDIP3 compared to both the control group and WT (Fig. 3E,F), suggesting that the G148V mutation disrupts normal splicing of POLDIP3. In addition, another CFTR exon 9 mini gene assay (Budini, Romano et al., 2015, Pinarbasi, Cagatay et al., 2018) was examined. The result clearly showed that co-expression with the reporter led to an enhanced signal of exon 9 exclusion for WT, while the exon 9 exclusion signal for the G148V mutation was significantly reduced, similar to the control group (Appendix Fig. S3C,D). In the signal of CFTR exon 9 not being cleaved, only WT demonstrated significant difference compared to other groups (Appendix Fig. S3C). These findings suggest that the G148V mutation impaired its function in the splicing process of CFTR exon 9. The findings suggest that the puncta formed by the TDP-43 G148V mutant may interfere with the normal splicing process within the cell nucleus. These cellular splicing defects are consistent with our earlier in vitro observations showing that the G148V mutation disrupts the DNA/RNA-binding ability of TDP-43. Accordingly, we propose that mutation-induced conformational changes lead to a loss of nucleic acid engagement, which in turn compromises TDP-43–dependent splicing regulation. In summary, under disease conditions, TDP-43 undergoes a biochemical redistribution, shifting from soluble pools toward detergent-insoluble assemblies. In line with this, the G148V variant displayed reduced soluble monomers and an enrichment of detergent-resistant high-molecular-weight species. Moreover, phosphorylation at pathological sites such as S409/410 was markedly elevated in the insoluble fraction, reinforcing the notion that G148V recapitulates ALS-linked biochemical hallmarks. Functionally, although nuclear aggregation of G148V partially alleviated cytotoxicity compared to wild type, splicing assays revealed impaired regulation of endogenous POLDIP3 exon 3 and diminished activity in the CFTR exon 9 minigene reporter, highlighting a disruption of RNA processing capacity.

### HSC70 colocalized with TDP-43 G148V puncta in the nucleus

Previous work showed that acetylation-mimicked TDP-43 organizes into nuclear anisosome structures, with HSP70 proteins concentrated in the core (Yu, Lu et al., 2021). These studies noted that when TDP-43 accumulates at the periphery of the anisosome, it encircles heat shock proteins such as HSPA1A, HSPA1L, HSPA6, and HSPA8, which are in the core of the anisosome (Yu et al., 2021). Furthermore, they found that the chaperone activity of HSP70 can sustain the anisosome structure of TDP-43 (Yu et al., 2021). Consequently, we investigated whether the puncta formed by TDP-43 G148V have any relationship with these heat shock proteins. Our experiments with HEK293T cells indicated that HSC70 (also known as HSPA8) predominantly localized in the cytoplasm of cells expressing control and wild-type TDP-43 (Fig. 5A). There are almost no HSC70-containing puncta in the nucleus. Conversely, in the cells expressing the TDP-43 G148V mutant, HSC70 accumulated in the nucleus, where its signal highly overlapped with the puncta formed by TDP-43 G148V (Fig. 5A). These observations were also observed in N2a cells (Appendix Fig. S4A). Quantitative analysis of the location of HSC70 demonstrated nearly 95% of HSC70-containing nuclear puncta in cells expressing TDP-43 G148V, but only ∼5% in the cells expressing wild-type TDP-43 (Fig. 5B). Analysis of the fluorescence spectrum profiles in HEK293T cells also revealed that while the signals from the control group and wild type did not resemble those of HSC70, the signals from the G148V variant and HSC70 were highly overlapping (Fig. 5C; Appendix Fig. S4D). Together, the results showed potential puncta formation between TDP-43 G148V and HSC70 within the nucleus. Further analysis on other HSPs including HSP70 and HSP90 found that HSP70 and HSP90 do not interact with the G148V mutant nuclear puncta (Appendix Fig. S4B,C). To further validate this phenomenon, we utilized Proximity Ligation Assay (PLA) to examine the HSC70 and TDP-43 variants We found the PLA signal can only be detected in cells expressing TDP-43 G148V, but no signal in the control group and wild type, confirming that the HSC70 and TDP-43 G148V form puncta together in the nucleus (Fig. 5D).

**Figure 5.**
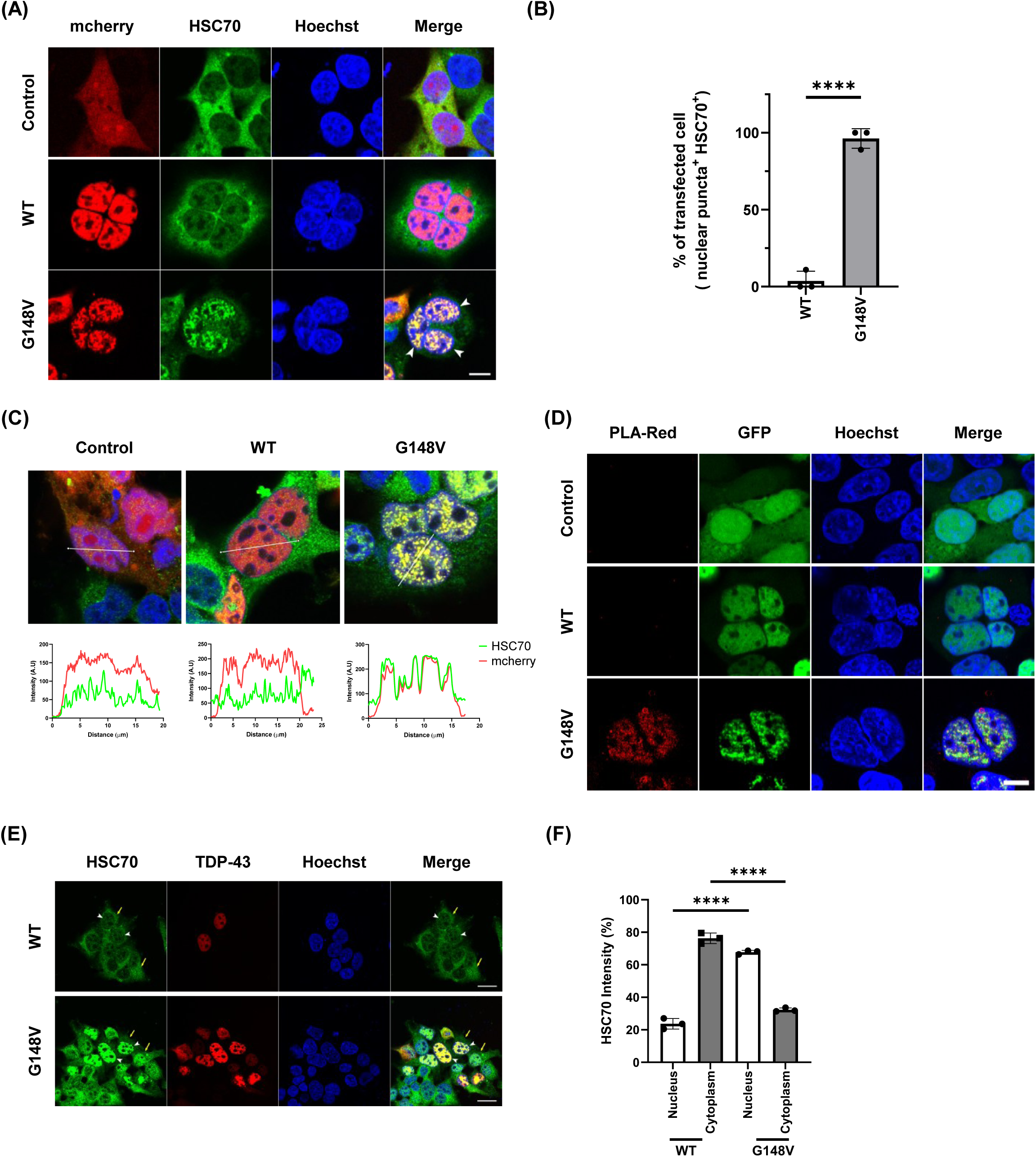
The TDP-43 G148V mutant affected the distribution of HSC70. (A) HSC70 co-localized with TDP-43 G148V puncta, but not TDP-43 WT, in the nucleus. Scale bar: 10 μm. (B) Quantitation of % of puncta with or without HSC70 in the cell nucleus. (n = 3 biological replicates) (C) Fluorescence profiles showed strong overlap between G148V and HSC70, suggesting their nuclear colocalization. (D) The proximity ligation assay between TDP-43 G148V puncta and HSC70. PLA signals were detected only in G148V-expressing cells, indicating nuclear colocalization with HSC70. Scale bar: 10 μm. (E) The distribution of HSC70 in the TDP-43 WT and G148V mutant expressed cells. (F) Quantification of HSC70 distribution in the nucleus and cytoplasm. HSC70 was mainly cytoplasmic in WT cells but redistributed to the nucleus in G148V cells. (n = 3 biological replicates) Data information: All data are presented as mean ± SD. In (E), the white arrow represents the nucleus, while the yellow arrow represents the cytoplasm. In (B)(F), n = ∼80 cells, ***P = 0.0006, ****P < 0.0001 (Student’s t-test).

HSC70 typically resides in the cytoplasm, performing chaperone functions such as chaperone-mediated autophagy (CMA) (Ormeno, Hormazabal et al., 2020, Valdor & Martinez-Vicente, 2024). Our results showed an alteration of the cellular distribution of HSC70 from cytoplasm to nucleus in the cells expressing G148V comparing to the wild-type. Quantitative analysis confirmed that, in wild-type cells, HSC70 diffused mostly in the cytoplasm with a small fraction in the nucleus (Fig. 5E,F). In G148V cells, HSC70 was primarily located in the nucleus, with only a minor presence in the cytoplasm (Fig. 5E,F). These might suggest that when TDP-43 G148V forms puncta in the nucleus, HSC70 might be affected and migrated from the cytoplasm to the nucleus.

### HSC70 maintained the solubility of TDP-43 G148V puncta in the nucleus to avoid cytotoxicity

To further examine the role of HSC70 in modulating G148V nuclear puncta, we employed an HSC70-specific siRNA to reduce HSC70 expression and examine changes in G148V puncta. We first titrated siRNA concentrations and observed a dose-dependent decrease in HSC70 protein levels (Appendix Fig. S5A,B). Based on these titration data, 100 nM siRNA was selected for subsequent experiments (Appendix Fig. S5A,B). In parallel experiments with and without HSC70 knockdown, HSC70 siRNA markedly reduced HSC70 signal relative to control, whereas no obvious morphological differences were observed between TDP-43 wild-type and G148V mutant cells (Appendix Fig. S5C,D), indicating that HSC70 depletion did not affect formation of G148V nuclear puncta. We next interrogated whether HSC70 knockdown altered the biophysical properties of G148V puncta. Fluorescence recovery after photobleaching (FRAP) experiments reproduced prior observations that G148V puncta display both liquid-like and solid-like behaviors (Fig. 6A,B). Quantification revealed that HSC70 depletion significantly increased the fraction of solid-like G148V puncta compared with control, with a concomitant reduction in the fraction of liquid-like puncta (Fig. 6C), suggesting that HSC70 contributed to maintaining the G148V puncta solubility. Prompted by these findings, we examined biochemical solubility and phosphorylation of TDP-43 G148V by biochemical fractionation and western blotting. Both representative blots and quantitative analysis demonstrated a pronounced increase—exceeding an order of magnitude—in phosphorylated, insoluble TDP-43 G148V following HSC70 knockdown, whereas wild-type TDP-43 was minimally affected (Fig. 6D,E). These results indicated that HSC70 not only preserves the solubility of G148V nuclear puncta but also modulates their phosphorylation state. Given that increased insolubility and phosphorylation are established pathological features, we hypothesized that HSC70 depletion would reduce the viability of cells expressing the G148V mutant. In the scramble control group, consistent with our previous observations, although both WT- and G148V-expressing cells showed a marked reduction in viability, G148V-expressing cells retained significantly higher viability than WT-expressing cells (Fig. 6F). Upon HSC70 knockdown, viability decreased significantly for both genotypes relative to control; however, G148V-expressing cells exhibited a markedly larger loss of viability (≥14% reduction) compared with a ∼3.5% reduction in wild-type cells (Fig. 6F). From the survival data, we concluded that HSC70 depletion disproportionately compromises the survival of cells expressing the G148V mutant. Overall, our results supported a model in which nuclear translocation of HSC70 mitigated G148V puncta toxicity by maintaining nuclear puncta solubility and limiting their phosphorylation.

**Figure 6.**
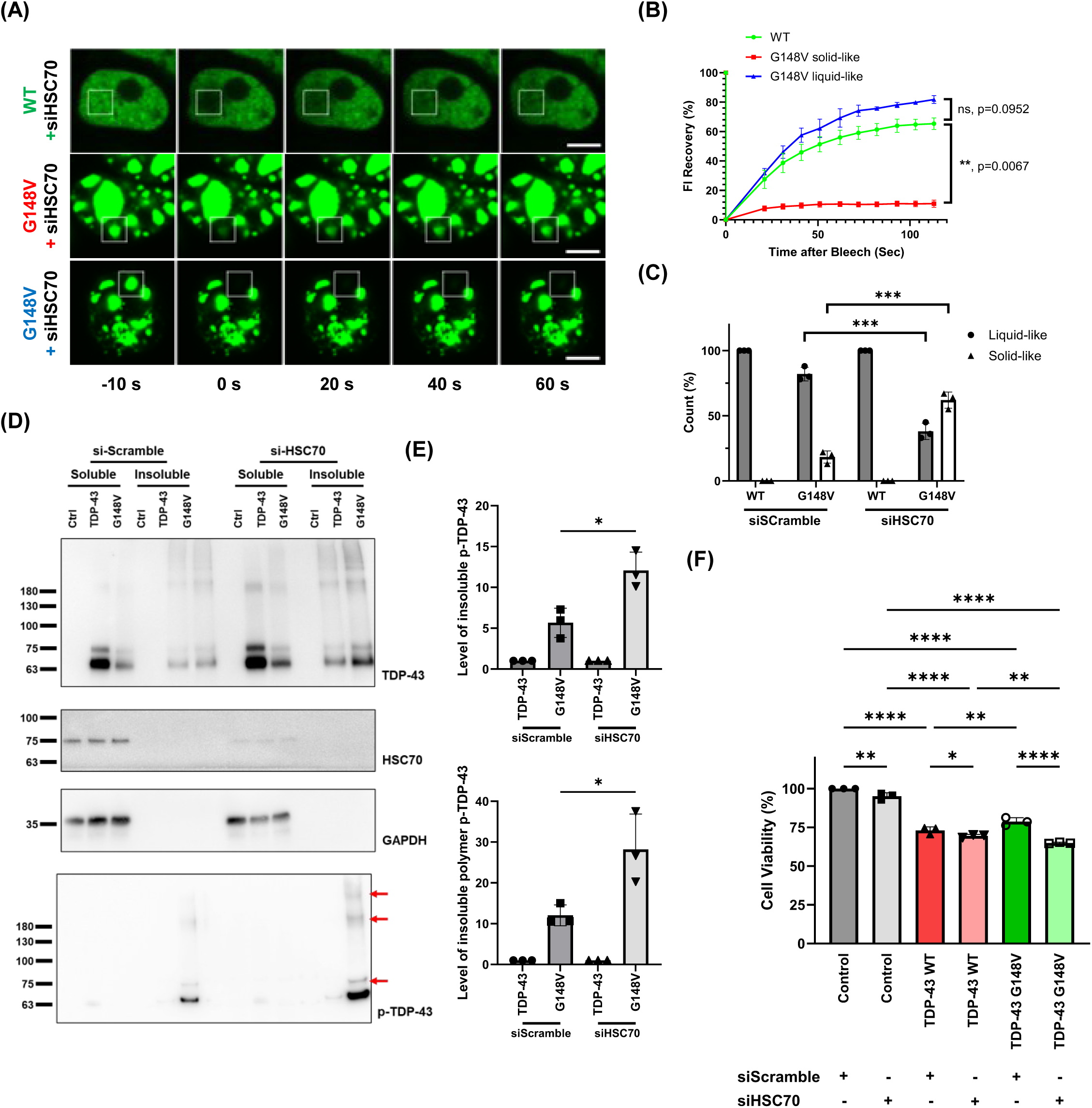
Knockdown of HSC70 influenced the characteristics and cell viability of G148V nuclear puncta. (A) The FRAP results of G148V with siHSC70. Scale bar: 5 μm. (B) The trend chart between WT and solid-like G148V puncta. (n = 3 biological replicates). (C) Quantitative result of FRAP with or without siHSC70. HSC70 depletion shifted G148V puncta from a liquid-like to a solid-like state. (n = 3 biological replicates). (D) The expression level of TDP-43 G148V with scramble siRNA or HSC70-specific siRNA. (E) Quantitative results of p-G148V mutant with scramble siRNA or HSC70-specific siRNA. HSC70 knockdown markedly increased phosphorylated, insoluble G148V, with minimal effect on WT. (n = 3 biological replicates). (F) CCK-8 assay demonstrated that HSC70 knockdown reduced viability in both WT- and G148V-expressing cells, with a greater loss in G148V cells. (n = 3 biological replicates) Data information: All data are presented as mean ± SD. In (B), n = ∼20 cells. **P = 0.0067 (Two-way Anova). In (C), ***P = 0.0007 (liquid-like) and 0.0006 (solid-like) (Student’s t-test). In (D), the upper and lower red arrows indicate polymer and monomer, respectively. In (E), *P = 0.0179 and 0.0371 (Student’s t-test). In (F), *P < 0.05, **P < 0.01, ***P < 0.001, ****P < 0.0001 (Two-way Anova with Tukey’s multiple comparisons test)

### G148V mutation drove nuclear puncta formation and HSC70 mislocalization as a model for early ALS

Through a series of examinations, we identified the G148V mutant as a promising candidate for establishing a cellular model to study early-stage amyotrophic lateral sclerosis (ALS). Using this mutant as a biological model, we further observed that HSC70 is associated with the dynamics of nuclear puncta and with cell viability. To validate whether this correlation is disease-related, we employed two previously reported pathogenic mutants: K181E, identified in ALS patients, and K263E, identified in FTLD patients (Chen et al., 2019). Both mutants have been described in the literature to form nuclear puncta when expressed in HEK293T cells, with a frequency of approximately 30% (Chen et al., 2019). In our experiments, overexpression of K181E, K263E, and G148V each led to the formation of nuclear puncta (Fig. 7A). Quantitative analysis confirmed that, consistent with prior reports, K181E and K263E formed nuclear puncta in ∼25% of cells, whereas the G148V mutant exhibited a strikingly higher frequency, with >80% of cells displaying nuclear puncta (Fig. 7B). Interestingly, when K181E and K263E formed nuclear puncta, HSC70 distribution was also altered, re-localizing from the cytoplasm into the nucleus (Fig. 7A). Although the extent of HSC70 mis-localization was lower than that observed with G148V, it was still significantly different from wild-type (Fig. 7C). To further substantiate these findings, we performed parallel experiments in N2a cells, which yielded results consistent with those in HEK293T cells: the G148V mutant displayed a markedly higher propensity to form nuclear puncta compared to K181E and K263E, and HSC70 redistribution accompanied puncta formation (Fig. 7D-F). Importantly, HSC70 mislocalization was observed not only in the G148V mutant but also in the known human-derived mutants, underscoring a shared pathological feature. Taking together, these results demonstrate that, compared with other mutants, G148V exhibits a substantially higher probability of nuclear puncta formation while also inducing HSC70 mislocalization, similar to known ALS- and FTLD-associated mutations. These findings highlight G148V as a highly promising cellular model for investigating early ALS pathogenesis.

**Figure 7.**
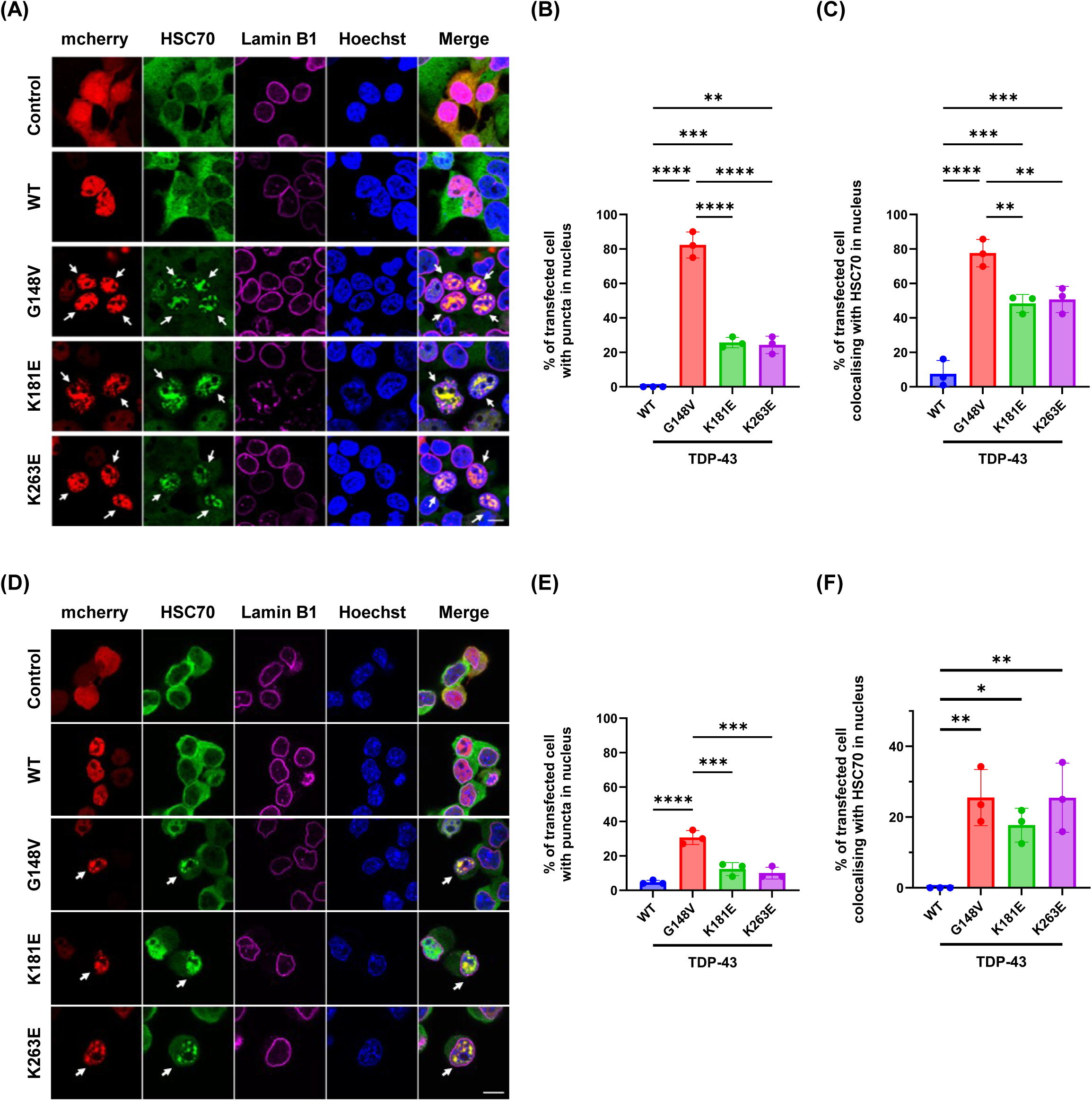
Both K181E and K263E, similar to G148V, exhibited aberrant localization of HSC70. (A) HSC70 co-localized with TDP-43 K181E and K263E puncta in the HEK293T nucleus. Scale bar: 20 μm. (B) Quantitative result of puncta in the transfected cell nucleus. (C) Quantitative result of puncta with HSC70 in the transfected cell nucleus. (D) HSC70 co-localized with TDP-43 K181E and K263E puncta in the N2a nucleus. Scale bar: 20 μm. (E) Quantitative result of puncta in the transfected cell nucleus. (F) Quantitative result of puncta with HSC70 in the transfected cell nucleus. Data information: All data are presented as mean ± SD. In (B)(C)(E)(F), n = ∼80 cells, *P < 0.05, **P < 0.01, ***P < 0.001, ****P < 0.0001 (One-way Anova with Tukey’s multiple comparisons test)

## Discussion

In this study, we employed a single RRM-adjacent mutation, G148V, and demonstrated that it is a potent driver of nuclear TDP-43 puncta. These puncta can be recapitulated in mammalian cells and reproduce pathological features known from ALS (Huang et al., 2024, Keating et al., 2023, Scherer, Maurel et al., 2024, Spence et al., 2024), including solid-like material properties, formation of insoluble species, hyperphosphorylation, and reduced cell viability. Recombinant protein analyses show that the G148V substitution alters TDP-43 conformation and abolishes its RNA/DNA binding, thereby impairing normal splicing function. Integrating the G148V data with prior literature, we reveal a modulatory role for HSC70 in nuclear TDP-43 assemblies: in mammalian cells, HSC70 is recruited from the cytoplasm into nuclear TDP-43 puncta, altering its cellular distribution. Using specific siRNA to deplete HSC70, we found that G148V more readily forms solid-like nuclear puncta with increased phosphorylation and a marked decrease in cell survival. These findings suggest that nuclear HSC70 helps maintain puncta fluidity and restrain pathological phosphorylation, thereby mitigating puncta-associated cytotoxicity. Finally, by comparing G148V with other disease-associated human variants (K181E and K263E) (Chen et al., 2019, Kovacs, Murrell et al., 2009), we validate G148V as a promising cell-biological model for studying early events in ALS pathogenesis.

We attempted to use AlphaFold-based modeling to evaluate the potential structural effects of the G148V substitution on TDP-43. Although the full-length protein predictions did not reveal marked changes in the overall TDP-43 conformation, modeling of the RRM1 and RRM2 region with DNA suggested a difference between WT and G148V. WT appeared to interact with the DNA in a more extended conformation, whereas G148V appeared to adopt a more compact state and contacted only a limited portion of the probe (Appendix Fig. S6A). Consistent with these observations, our biophysical measurements also revealed detectable structural consequences. Far-UV circular dichroism and intrinsic fluorescence both point to alterations in secondary and tertiary structure: the G148V variant shows an increased α-helical signal and reduced solvent exposure of aromatic side chains, consistent with a modest compaction of local structure. Notably, AlphaFold models did not predict changes in the number of hydrogen bonds within RRM1, suggesting that the mutation perturbs local packing or dynamics rather than breaking core hydrogen-bonding networks. Given that residues F147 and F149 flank G148 and are known contributors to nucleic-acid recognition (Kuo et al., 2014), these structural shifts led us to hypothesize an effect on nucleic-acid engagement. Indeed, anisotropy assays demonstrate that G148V abolishes binding to TG single-stranded probes (Chang et al., 2012, Kuo et al., 2014), confirming a loss of DNA/RNA affinity. Loss of nucleic-acid interactions is expected to reduce the solvation and multivalent contacts that normally stabilize dynamic, RNA-bound TDP-43 assemblies; consequently, the protein becomes more prone to self-association (Maharana, Wang et al., 2018). We therefore propose that the G148V substitution undermines TDP-43’s nucleic-acid chaperoning function, lowering its solubility and promoting the nucleation of nuclear puncta. Over time, these assemblies can mature toward more solid-like states, increasing phosphorylation and insolubility and producing downstream cellular effects.

In recent years, a growing body of work has implicated aberrant nuclear TDP-43 function as an early hallmark of ALS (Carmen-Orozco, Tsao et al., 2023, Decker et al., 2025, Huang et al., 2024, Spence et al., 2024). Clinical analyses published in 2024 reported nuclear TDP-43 accumulation in brain tissue from presymptomatic individuals, accompanied by disrupted splicing of STMN2 (Spence et al., 2024). Another study induced nuclear TDP-43 condensation using chemical stressors and likewise documented widespread perturbations in known splicing targets (Huang et al., 2024). Together, these reports support a model in which nuclear aggregation of TDP-43 precedes the canonical cytoplasmic mislocalization and inclusion formation that ultimately culminate in neuronal loss. Consistent with these observations, we identified a single RRM-proximal substitution, G148V, that reproducibly drives nuclear TDP-43 puncta. Expression of G148V produces nuclear puncta at high penetrance (>80%) across multiple cell types without the need for exogenous stress, a frequency substantially greater than that observed for the human-derived variants K181E and K263E (Chen et al., 2019, Murrell et al., 2009). The G148V puncta display multiple ALS-associated biochemical and biophysical signatures, including pronounced phosphorylation at S409/S410, formation of SDS-resistant insoluble species, and a transition toward solid-like liquid properties. Functionally, these nuclear assemblies impair normal pre-mRNA processing, increasing cryptic exon inclusion and leading to reduced cell viability. Taken together, our data indicate that G148V models several early, disease-relevant features of TDP-43 proteinopathy and therefore represents a useful cellular system for probing initial pathogenic events. Although we have validated key phenotypes in the N2a neuroblastoma line, this model does not fully recapitulate mature motor neuron biology. Accordingly, we plan to extend these studies into primary neurons and iPSC-derived motor neurons to confirm the generality of the observed effects and to evaluate additional cryptic-exon candidates such as STMN2 and UNC13A (Huang et al., 2024).

HSC70 participates in multiple cellular regulatory pathways; it is best characterized for roles in clathrin-mediated endocytosis and chaperone-mediated autophagy (Ciechanover & Kwon, 2017, Rai, Kennedy et al., 2021). An earlier paper showed that HSC70 directly participates in nuclear import by binding nuclear localization signals (NLSs) and escorting karyophilic proteins to the nuclear pore. Antibodies against HSC70 specifically blocked the mediated nuclear import of nucleoplasmin and SV40 T-antigen NLS-bearing cargo, demonstrating that HSC70 acts as a cytoplasmic NLS-binding factor through the nuclear pore complex. (Imamoto, Matsuoka et al., 1992). CMA driven by HSC70 has been linked to the clearance of aberrant cytoplasmic TDP-43 assemblies (Huang, Bose et al., 2014, Ormeno et al., 2020). Previous literatures have demonstrated that there is an directly interaction between HSC70 and TDP-43 in vitro and in vivo (Huang, Bose et al., Francois-Moutal, Scott et al., 2022). Together, suggesting that the aggregate-prone form of TDP-43 is able to interact with Hsc70, to co-localize with Lamp2A, and to up-regulate the levels of these molecular components of CMA. Additionally, in ALS patients’ peripheral blood mononuclear cells (PBMCs) from peripheral blood of patients, HSC70 expression has been reported to decline, suggesting that reduced chaperone capacity may impair the removal of dysfunctional TDP-43 and contribute (Arosio, Cristofani et al., 2020). Although HSC70 functions, such as chaperone-mediated autophagy (CMA), are typically cytosolic, the small puncta induced by the TDP-43 G148V mutation elicit atypical HSC70 responses. Notably, in cells expressing the G148V TDP-43 variant, we observed an altered subcellular distribution of HSC70. Rather than remaining predominantly cytosolic, HSC70 translocated into the nucleus and co-accumulates with the small nuclear puncta formed by G148V. By contrast, the HSP70 and HSP90 did not show comparable relocalization, indicating a selective association of HSC70 with these nuclear assemblies. Functionally, reduction of HSC70 levels increased the proportion of solid-like nuclear puncta, elevated the abundance of phosphorylated, detergent-insoluble TDP-43 species, and exacerbated loss of cell viability. These findings are consistent with a model in which HSC70 acts as a regulator of both the material properties and the post-translational modification state of TDP-43 assemblies, mitigating toxicity by preserving liquidity and limiting pathological phosphorylation.

Our current results, however, describe end-points and do not yet resolve the intermediate steps that connect HSC70 relocalization to altered TDP-43 biochemistry. Several key questions remain. First, it is unclear whether HSC70 associates with TDP-43 before nuclear entry or is recruited after TDP-43 has accumulated in the nucleus. Given prior evidence that HSC70 can bind TDP-43 and may participate in nucleo-cytoplasmic transport, one plausible hypothesis is that HSC70 escorts TDP-43 into the nucleus following a cytosolic encounter; in the case of G148V, altered client dynamics could trap HSC70 within the nucleus, which paradoxically attenuates acute toxicity by stabilizing nascent puncta. To test this, in vitro binding assays using purified proteins could determine whether G148V alters HSC70–TDP-43 affinity, and live-cell imaging combined with perturbation of nuclear transport pathways could clarify the temporal and spatial sequence of their interaction. Second, the mechanism by which HSC70 depletion increases phosphorylated, insoluble TDP-43 is unresolved. One possibility is that HSC70 binding sterically occludes kinase or phosphatase docking sites on TDP-43; loss of HSC70 would therefore expose these sites and shift the balance toward hyperphosphorylation. Pharmacological or genetic modulation of candidate kinases and phosphatases could distinguish whether altered enzyme access underlies the phosphorylation changes. Third, the functional significance of HSC70 nuclear entry warrants further study because canonical CMA operates in the cytosol. One hypothesis is that HSC70 attempts to extract aberrant nuclear TDP-43 for cytosolic degradation but becomes sequestered within large or high-affinity nuclear assemblies. Alternatively, HSC70 might participate in a nuclear protein quality-control pathway distinct from classical CMA; recent work describing nucleolus-associated clearance mechanisms raises the possibility that HSC70 contributes to such nuclear proteostasis systems (Brunello, Polanowska et al., 2025). Addressing these possibilities will require experiments that probe nuclear export competence, measure the dynamics of HSC70 retention, and test involvement of known nuclear quality-control factors. Collectively, these lines of inquiry will clarify whether HSC70’s nuclear relocalization reflects an adaptive clearance attempt, a maladaptive sequestration, or a combination of both, and will define how chaperone dynamics influence the transition from reversible condensates to pathological, insoluble TDP-43 assemblies.

In this study, we demonstrate a direct connection between an RRM-proximal structural perturbation (G148V) and the formation of nuclear TDP-43 condensates, a process that is mitigated by the molecular chaperone HSC70. Nuclear HSC70 appears to preserve the fluidity of these assemblies and to limit phosphorylation-driven transitions to insoluble states, identifying it as a key regulator of early TDP-43 proteinopathy. Importantly, G148V induces robust and reproducible nuclear puncta formation in the absence of exogenous chemical stressors or inhibitors, providing an efficient cellular platform for the mechanistic dissection of nuclear pathology and for evaluating interventions that target chaperone function or restore TDP-43 RNA-binding activity.

## Methods

### Plasmids

Full length of TDP-43 in the pEGFP-C3 and pCMV-Tag2B plasmids are gifts from Dr. Tu Pang-Hsien, Institute of Biomedical Sciences, Academia Sinica, Taiwan and pAG426-TDP-43 plasmids are gifts from Dr. Aaron Gitler, Stanford University, California, USA. For the recombinant protein production, His-tagged TDP-43 in pET14b was used as described in the previous study (Fang YS et al. 2014). TDP-43 G148V, K181E, and K263E mutant in all plasmids were generated by site-directed mutagenesis with specific primers. The mcherry sequence was amplified and introduced into the N-terminal of pCMB-Tag2B using Gibson assembly Master Mix assembly (E2611, NEB) to form pCMV-Tag2B-mcherry variants. All plasmids were confirmed by Sanger DNA sequencing.

### Protein purification

The N-terminal His-tagged TDP-43 WT and G148V mutant were transformed and over-expressed in the Rosetta 2 E. coli strain (Novagen, Merck KGaA). The cells were cultured first in 100 ml LB containing ampicillin (100 mg/ml, Amp) and chloramphenicol (34 mg/ml, Cam) at 200 rpm, at 37 °C overnight. Then, subculturing to large scale (1 L) for 4-6 hours to reach OD600 = 0.6-0.8 and add Isopropyl β-D-1-thiogalactopyranoside (IPTG) at 25 °C overnight. The next day, the culture was centrifuged at 5,000 rpm, 4 °C for 15 minutes, discarded the supernatant, resuspended in 50 ml lysis buffer, and incubated at −20 °C overnight. After, added 100 μL of 10 mg/mL of DNAase I, 100 μL of 10 mg/mL of RNAase A, 1 mM of DTT (50 μL), and 100 μL proteinase cocktail inhibitors III, lysed the cell with High Pressure Homogenizer, centrifuge at 15,000 rpm, 4 °C for 30 minutes, and discarded the supernatant. Resuspended the pellet with Buffer A (50 mM Tris, 200 mM NaCl, 25 mM imidazole, and 6 M Urea, pH 8) and kept at 4 °C for overnight. The next day, centrifuged at 15,000 rpm at 4 °C for 30 minutes and collected the supernatant. The 0.2 μm syringe filter was used to remove debris and transferred into GE50 ml Superloop to purify by HisTrap™ HP Prepacked Columns (GE Healthcare, Cytiva) with Buffer B (50 mM Tris, 200 mM NaCl, 500 mM imidazole and 6 M Urea, pH 8). The protein was analyzed by SDS-PAGE with Coomassie Blue staining.

Superdex-200 10/300 GL analytical gel-filtration column (GE Healthcare Bio-Sciences AB) was performed with Buffer A. The recombinant protein was concentrated by Amicon® Ultra Centrifugal Filter, 30 kDa MWCO (Millipore) into 1 ml in Buffer A and injected into Superdex-200 column. Samples were collected in 100 μl volume by fraction collector. Again, the sample fractions were analyzed by SDS-PAGE with Coomassie Blue staining. All TDP-43 protein was kept at −20 °C before dialysis. For experiments, the TDP-43 protein was loaded into the Spectra/Por 6 Dialysis Tubing (Repligen) and dialyzed in the refolding buffer (10 mM Tris, pH 8.0). After 4 hours at 4 °C, refresh the buffer and dialyze it overnight. The next day, the concentration of collected protein was measured the absorption at 280 nm by DU800 ultraviolet−visible spectrophotometer (Beckman Coulter, Inc) and quantified with the extinction coefficient of 44,920 cm^−1^M^−1^ according to the equation described by (Pace, Vajdos et al., 1995).

### Far-UV circular dichroism

The refolded TDP-43 protein was dialyzed and diluted with refolding buffer to achieve a working concentration of 3 μM. Far-UV circular dichroism (CD) spectra were acquired at room temperature using the protein solution in a circular quartz cell (Hellma, Forest Hills, NY, USA) with a Jasco J-815 spectropolarimeter (Jasco Inc., Easton, MD, USA) having a 1-mm pathlength. Spectra were recorded within the range of 250 to 180 nm and corrected by subtracting the buffer background. The spectra data were analyzed using K2D3 online software (Louis-Jeune C, et al. 2012).

### Intrinsic fluorescence spectroscopy

Under excitation at 280 and 295 nm, the intrinsic fluorescence of 3 μM TDP-43 was collected in the range of 290 to 450 nm and 300 to 450. All experiments were performed using a FluoroMax-Plus fluorescence spectrophotometer (Horiba Scientific).

### Fluorescence anisotropy

TDP-43 WT and G148V, was diluted to 0.25, 0.5, 0.75, 1, 1.25, 1.5, and 1.75 μM, mixed with the 10 nM DNA probes containing 15 TG dinucleotide (TG)15 or 20 TT dinucleotide (TT)20, and incubated at room temperature for 30 minutes. Afterwards, the anisotropy of all the mixed samples was loaded into the Ultra-Micro Fluorescence Cell (Hellma, Forest Hills, NY, USA). Fluorescence signals were excited at 490 nm and emissions were collected at 525 nm (Horiba Scientific). The (TG)15 and (TT)20 probes were labeled by Alexa-488 at 5’ terminus (GENOMICS).

### Cell culture and transfection

HEK293T and N2a were selected and cultured in the Dulbecco’s modified Eagle medium (DMEM, Gibco) containing 10% fetal bovine serum (Gibco), and maintained at 37 °C, 5% CO_2_. The cells were seeded and incubated for at least 24 hours before transfection. Lipofectamine 3000 transfection reagent (Invitrogen) was used to deliver the plasmids into cells. For knocking down, the siRNA for HSPA8/HSC70 (Santa Cruz) and control siRNA (Santa Cruz) were co-transfected with other plasmids. After transfection for 48 hours, the cells were collected for further analysis.

### Immunocytochemistry

Cells were seeded on the poly-D-lysine-coated coverslip and incubated for 24 hours at 37 °C, 5% CO_2_. After transfection for 48 hours, cells were fixed with 4% paraformaldehyde for 30 minutes and wash with washing buffer for 3 times, 5 minutes. Cells were treated with 0.5% Triton X-100 for 10 minutes to lose the membrane. After washing, cells were blocked with 3 % BSA for 1 hour at RT, followed by incubating with primary antibody in blocking buffer for another 1 hour at RT. Then, cells were washed and incubated with secondary antibody for 1 hour at RT. Subsequently, Hoechst 33258 (H3569, Invitrogen) in the PBS buffer was treated with cells for 10 minutes at RT. Finally, cells were mounted by prolong gold antifade (Invitrogen) and analyzed by SP8 LIGHTNING Confocal Microscope (Leica) with 63X oil-immersion objective (HC PL APO 63x/1,40 OIL CS2). All images and intensity profile were analyzed using FIJI ImageJ and Prism.

### Fluorescence recovery after photobleaching

Cells were seeded on the poly-D-lysine-coated coverslip and incubated for 24 hours at 37 °C, 5% CO_2_. After transfection for 48 hours, cells were analyzed by SP8 LIGHTNING Confocal Microscope (Leica) with 63X oil-immersion objective (HC PL APO 63x/1,40 OIL CS2). Green fluorescence bleaching was performed using a 488 nm wavelength argon laser at 100% intensity with 10 repetitions. Fluorescence recovery was monitored at 10-second intervals for a duration of two minutes. All images and intensity profile were analyzed using FIJI ImageJ and Prism.

### Western blotting analysis

For cell culture, cells were seeded on the poly-D-lysine-coated coverslip and incubated for 24 hours at 37 °C, 5% CO_2_. After transfection for 48 hours, cells were first washed with cold 4 °C PBS and collected by Protein lysis buffer (Goal Bio). After incubating for 10 minutes, samples were centrifuged at 15,000 rpm, 4 °C for 15 minutes and collected the supernatant for soluble fraction. Pellet samples were dissolved with Urea buffer (100 mM Tris, 20% glycerol, 4% SDS, and 8 M Urea, pH 6.8) by sonication. Samples were centrifuged at 15,000 rpm, RT for 15 minutes and collected as insoluble fraction. All samples were stored at −80 °C. The concentration of protein was determined by Pierce™ BCA Protein Assay Kits (Thermo Scientific). Sodium dodecyl sulfate-polyacrylamide gel electrophoresis (SDS-PAGE) was utilized to separate the protein and transferred to nitrocellulose membranes to immunoblot. All images were analyzed using FIJI ImageJ and Prism.

### Cell viability and ROS assay

Cells were seeded on the 96 well plate and incubated for 24 hours at 37 °C, 5% CO_2_. The flag-only plasmids were used in these experiments. After transfection for 48 hours, WST8 (2098, Cyrusbioscience) and general oxidative stress indicator (C6827, Invitrogen) were added into the cell samples and incubated for 1 hour at 37 °C, 5% CO_2_. After incubation, CCK-8 assay was analyzed by measuring the absorbance at 450 nm. Fluorescence signals of ROS were excited at 490 nm and emissions were collected at 525 nm. All results were analyzed using Prism.

### RNA extraction and RT-PCR

Cells were seeded on the 6 well plate and incubated for 24 hours at 37 °C, 5% CO_2_. After transfection for 48 hours, cells were collected and washed with cold PBS. RNeasy Mini Kit (Qiagen) was utilized to extract the RNA from cell samples. After normalizing all RNA samples to 500 ng, SuperScript™ IV Reverse Transcriptase (18090050, Invitrogen) was used. cDNA from reverse transcription was examined by PCR with POLDIP3 (Tollervey et al., 2011) and CFTR primers. For CFTR mini gene assay, cells needed to be co-transfected with pCDNA3.1 (+)-CFTR and TDP-43 variants in pCMV-Tag2B. All the PCR products were analyzed in 2% agarose gel and imaged by Bio-Rad Molecular Imager Gel Doc XR+ Imaging System Lab. All images were analyzed using FIJI ImageJ and Prism.

### Proximity ligation assay

Cells were seeded on the poly-D-lysine-coated coverslip and incubated for 24 hours at 37 °C, 5% CO_2_. After transfection for 48 hours, cells were fixed with 4% paraformaldehyde for 30 minutes and wash with washing buffer for 3 times, 5 minutes. Cells were treated with 0.5% Triton X-100 for 10 minutes to lose the membrane. After washing, the protocol in NaveniFlex Cell Red kit (60025, Navinci) was followed. Finally, Hoechst 33258 (H3569, Invitrogen) in the PBS buffer was treated with 10 minutes at RT and the samples were mounted by prolong gold antifade (Invitrogen) and analyzed by SP8 LIGHTNING Confocal Microscope (Leica) with 63X oil-immersion objective (HC PL APO 63x/1,40 OIL CS2). All images were analyzed using FIJI ImageJ.

## Supporting information

Reagent and Tools

## Data availability

This study includes no data deposited in external repositories.

## Acknowledgements

We would like to thank the confocal microscopy facility in the Genomics Research Center, Academia Sinica, for fluorescence imaging. We thank Academia Sinica Advanced Optics Microscope Core Facility for microscope imaging technical support. The core facility is funded by Academia Sinica Core Facility and Innovative Instrument Project (AS-CFII-114-A3). The research was supported by Academia Sinica, Taiwan (AS-IA-112-L07) and National Science and Technology Council, Taiwan (NSTC 115-2320-B-001 -019 -MY20)

## Author information

### Contributions

**Jeng Jung Wu:** Conceptualization; Data curation; Investigation; Methodology; Supervision; Writing—original draft. **Shin-Yu Fan:** Investigation. **Tien-Hsien Chang:** Supervision. **Yun-Ru Chen:** Conceptualization; Supervision; Funding acquisition; Writing—original draft; Project administration; Writing—review and editing.

### Ethics declarations

The authors declare no competing interests.

