## Appendix Figure for "HSC70 prevents TDP-43 nuclear puncta formation and toxicity in a novel nuclear puncta cell model for ALS"

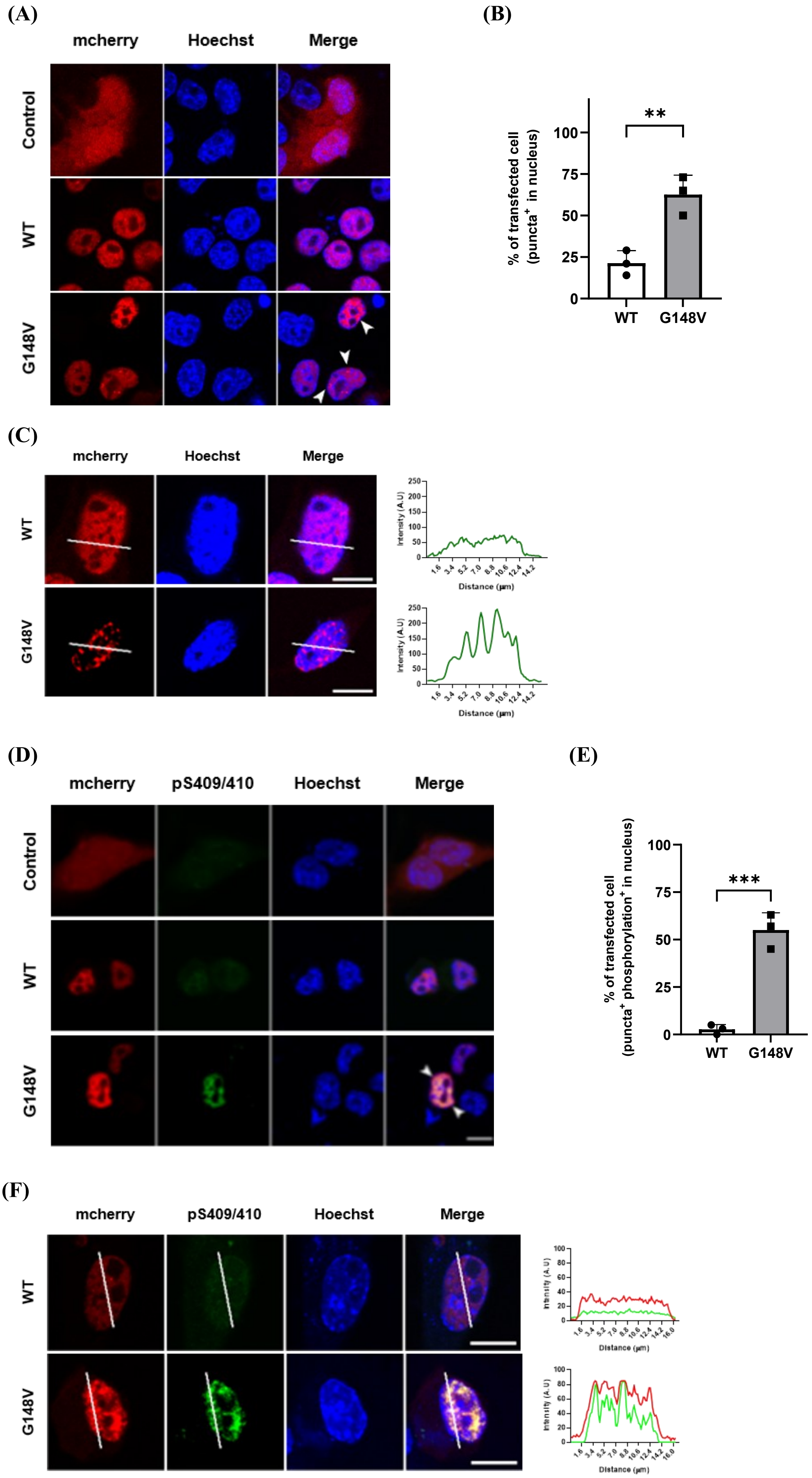

(A)
mcherry
Hoechst
Merge
Control
WT
G148V
(B)
% of transfected cell
(puncta+ in nucleus)
**
100
75
50
25
0
WT
G148V
(C)
mcherry
Hoechst
Merge
WT
G148V
Intensity (A.U)
Distance (μm)
(D)
mcherry
pS409/410
Hoechst
Merge
Control
WT
G148V
(E)
% of transfected cell
(puncta+ phosphorylation+ in nucleus)
***
100
75
50
25
0
WT
G148V
(F)
mcherry
pS409/410
Hoechst
Merge
WT
G148V
Intensity (A.U)
Distance (μm)

**Appendix Figure S1. The G148V mutation of TDP-43 formed nuclear puncta in N2a cells.**

(A) TDP-43 G148V formed nuclear puncta while WT demonstrated a more diffused form in the nucleus of N2a cells. Scale bar: 10 μm. (B) Quantitative result showed a significant difference between WT and G148V (n = 3 biological replicates). (C) The representative cell images for panel A. The intensity peaks of G148V are more heterogenicity than WT in the nucleus. Scale bar: 10 μm. (D) Phosphorylation level increased in the TDP-43 G148V puncta of in the nucleus of N2a cells. Scale bar: 10 μm. (E) Quantitative result showed a significant difference between WT and G148V (n = 3 biological replicates). (F) The representative cell images for panel E. The intensity peaks of G148V and pS409/S410 overlapped in the nucleus. Scale bar: 10 μm.

Data information: All data are presented as mean ± SD. In (B)(E), n = ~80 cells, **P = 0.0067, ***P = 0.0007 (Student's t-test).

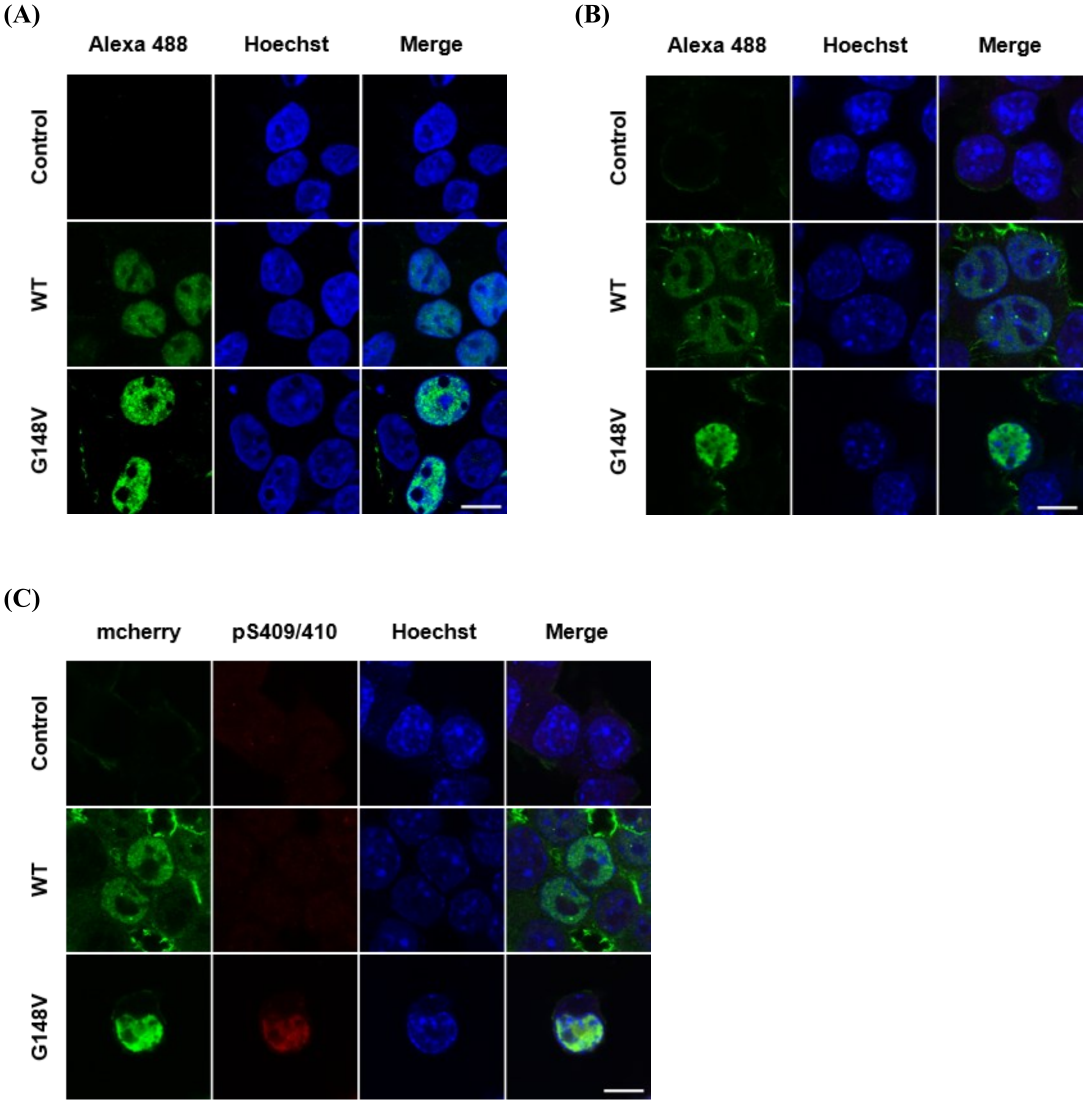


**Appendix Figure S2. The G148V mutation of flag-tagged TDP-43 also formed nuclear puncta in both HEK293T and N2a cells.**

(A) Flag-tagged TDP-43 G148V formed puncta while WT demonstrated a more diffused form in the nucleus of HEK293T cells. Scale bar: 10 μm. (B) Flag-tagged TDP-43 G148V formed puncta while WT demonstrated a more diffused form in the nucleus of N2a cells. Scale bar: 10 μm. (C) Phosphorylation level increased in flag-tagged TDP-43 G148V in the nucleus of N2a cells. Scale bar: 10 μm.

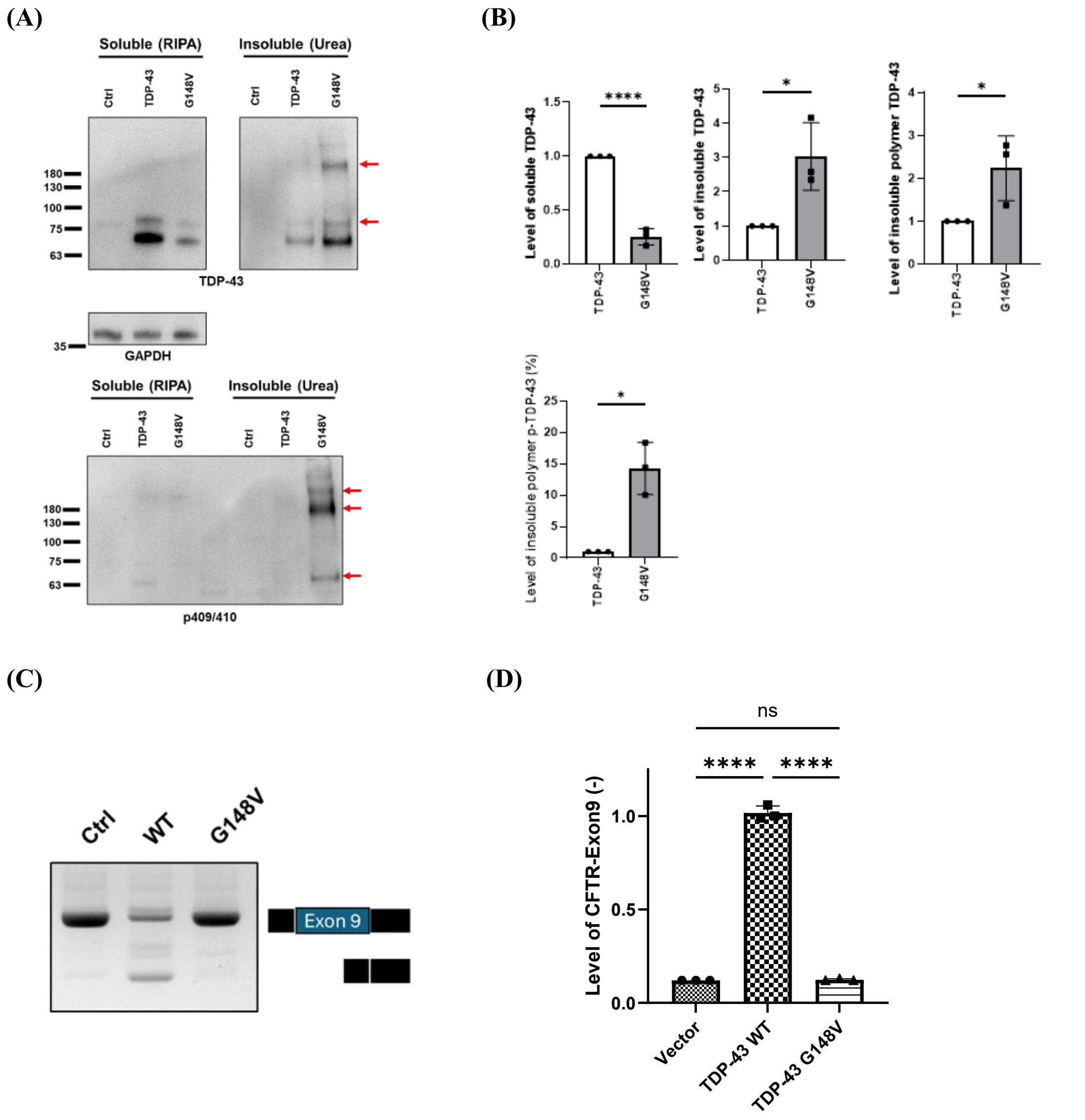


**Appendix Figure S3. The G148V mutation reduced cell viability but impaired CFTR splicing.**

(A) Expression levels of TDP-43 G148V in N2a cells. Soluble protein was extracted with RIPA buffer, and insoluble protein with urea buffer. (B) Quantification of the western blot result. G148V showed increased insoluble, hyperphosphorylated monomer and SDS-resistant HMW species compared with WT (n = 3 biological replicates). (C)

Representative RT-PCR of the CFTR exon 9 mini-gene assay. (D) Quantification of CFTR exon 9(-) signal (n = 3 biological replicates).

Data information: All data are presented as mean ± SD. In (A), the upper and lower red arrows indicate polymer and monomer, respectively. In (B), *P < 0.05, ****P < 0.0001 (Student’s t-test). In (D, E), *P = 0.0294, **P < 0.0001 (One-way Anova with Tukey's multiple comparisons test).

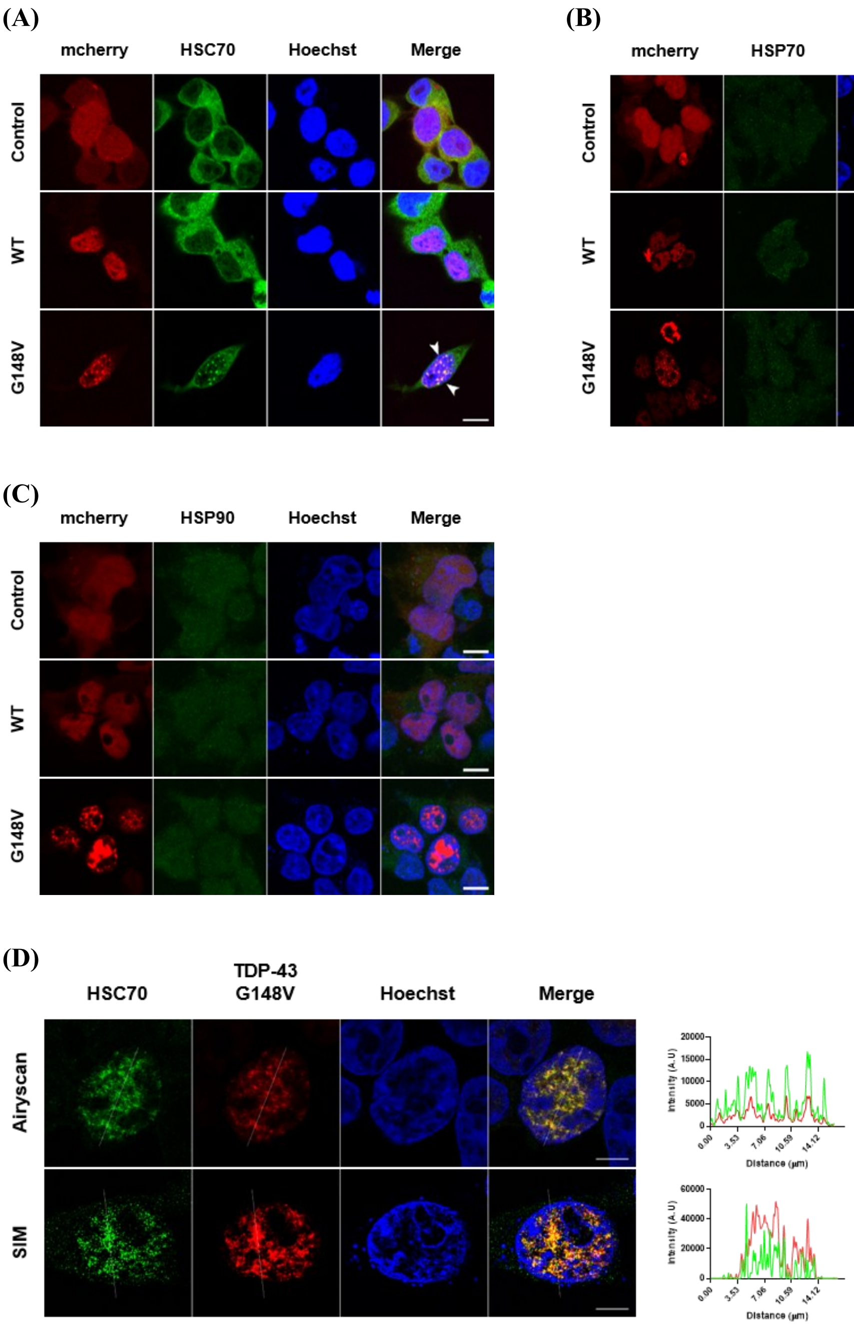


**Appendix Figure S4. HSC70, but not HSP70 and HSP90, co-localized with TDP-43 G148V puncta.**

(A) HSC70 co-localized with TDP-43 G148V puncta, but not TDP-43 WT, in the nucleus. Scale bar: 10 μm. The colocalization was indicated by yellow arrows. (B, C)

HSP70 (B) and HSP90 (C) didn’t co-localize with TDP-43 G148V puncta in the nucleus. Scale bar: 10 μm. (D) The images of G148V and HSC70 in airyscan and SIM microscopy. Scale bar: 10 μm.

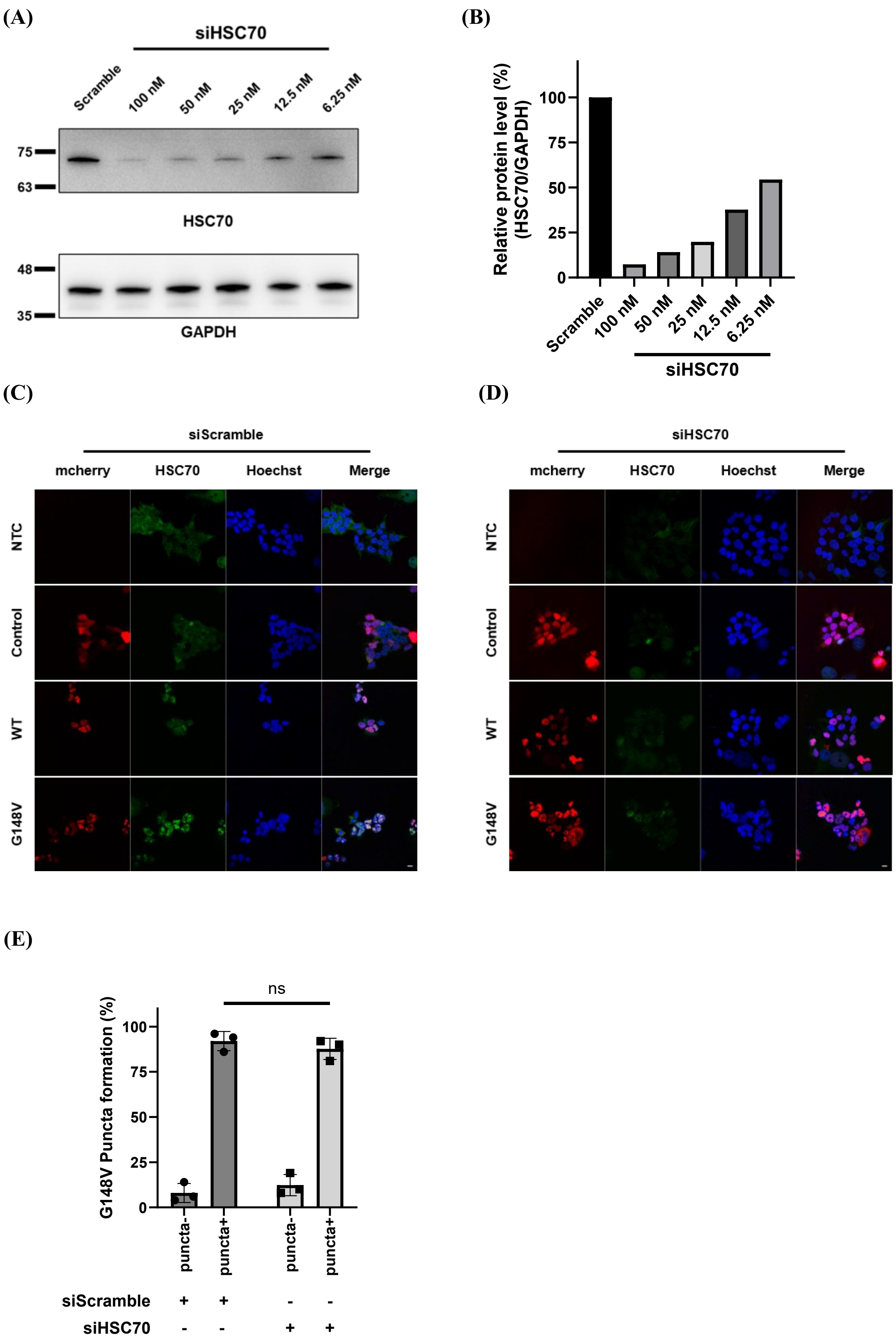
(A)
siHSC70
Scramble
100 nM
50 nM
25 nM
12.5 nM
6.25 nM
75
63
HSC70
48
35
GAPDH
(B)
Relative protein level (%)
(HSC70/GAPDH)
100
75
50
25
0
Scramble
100 nM
50 nM
25 nM
12.5 nM
6.25 nM
siHSC70
(C)
siScramble
mcherry
HSC70
Hoechst
Merge
NTC
Control
WT
G148V
(D)
siHSC70
mcherry
HSC70
Hoechst
Merge
NTC
Control
WT
G148V
(E)
ns
G148V Puncta formation (%)
100
75
50
25
0
puncta-
puncta+
puncta-
puncta+
siScramble + + - -
siHSC70 - - + +

**Appendix Figure S5. HSC70 knockdown reduced HSC70 levels without affecting G148V nuclear puncta.**

(A) The knockdown efficiency of HSC70. The titration result of HSC70-specific siRNA in HEK293T was shown. (B) The quantification of the Western blot in knockdown HSC70 shown in panel A. (C, D) The fluorescent images of TDP-43 WT and G148V with scramble siRNA (C) and HSC70-specific siRNA (D). Scale bar: 10 μm. (E) Quantitative result of the G148V transfected cells with the nuclear puncta with siScramble or siHSC70 (n = 3 biological replicates).

Data information: All data are presented as mean ± SD. In (E), n = ~80 cells, ns = non-significant (Student's t-test).

(A)

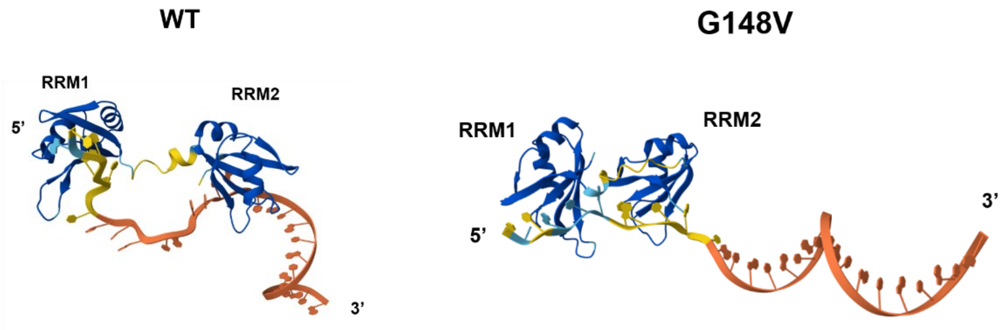


(B)

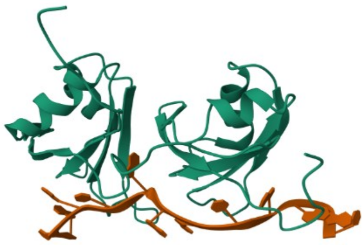

**Appendix Figure S6. The G148V mutation may lead to a relatively more compact conformation of the RRM1–RRM2 region.**

(A) The RRM1 and RRM2 domain with TG15 DNA probe insight structural prediction of TDP-43 WT (right) and G148V (left) from AlphaFold (https://alphafoldserver.com/). Compared with WT, the G148V mutation may induce a relatively more compact conformation. (B) The RRM1 and RRM2 domain with UG-rich RNA from NMR (PDB: 4B2S).
